# Structural model of the blood protein vitronectin bound to lipid and HDL

**DOI:** 10.1101/2025.11.01.685992

**Authors:** Kyungsoo Shin, William Brown, Ye Tian, Tata Gopinath, Andrey A Bobkov, Fabrizio Marinelli, Francesca M. Marassi

## Abstract

Vitronectin (Vn) is a multifunctional blood glycoprotein involved in cell adhesion and migration, blood coagulation, and inflammation. It is a component of the high-density lipoprotein (HDL) proteome also found associated with the calcified, lipid-rich, protein deposits that are a hallmark of age-related macular degeneration, Alzheimer’s disease, atherosclerosis and other aging-related diseases. Here we explored the molecular basis for lipid binding and HDL association by Vn. Using nuclear magnetic resonance (NMR) and all-atom molecular dynamics (MD) simulations, we identify a hydrophobic groove on the surface of the Vn hemopexin-like (HX) domain, that is capable of binding phospholipids. Key residues mediate the association of VnHX with phosphatidylcholine (PC) via hydrophobic and polar contacts. Lipid binding is guided by interactions with positively charged and hydrophobic residues that organize the lipids in a tail-to-tail bilayer-like arrangement within the groove. MD simulations of PC-bound VnHX combined with an MD structural model of discoidal HDL, show that VnHX associates with HDL in a lipid-mediated manner and at the exposed edge of the discoidal HDL lipoparticle where the terminal helices of the scaffolding protein ApoA1 flip away. Collectively, the data establish a comprehensive structural model for Vn association with PC and HDL. They expand the range of protein-lipid interactions observed within the structurally and functionally diverse collection of HDL particles.

## INTRODUCTION

High-density lipoprotein (HDL) is a heterogeneous collection of lipid-protein particles with fundamental activities in human health (1-4). The primary building blocks of HDL are the helical scaffolding protein ApoA1, phospholipids and unesterified cholesterol, but the HDL proteome and lipidome include more than 280 protein and 200 lipid species that endow it with a broad range of important functionalities, including lipid homeostasis, inflammation, innate immunity, and glucose control (5-9). HDL is best known for its protective activity against cardiovascular disease, with low levels of plasma HDL cholesterol and particle size correlating with elevated risk of cardiovascular diseases. This protective property stems from the interaction of nascent discoidal HDL with lecithin-cholesterol acyltransferase, which converts cholesterol to cholesteryl ester resulting in spherical αHDL that is absorbed into the liver during reverse cholesterol transport. Precisely how HDL adjusts its size and structure to accommodate specific levels of phospholipid, cholesterol and additional proteins in this process, is an important question for understanding its activities. Structural studies have provided fundamental information for both the nascent discoidal and mature spherical HDL particles (10), and structural data are emerging for lecithin-cholesterol acyltransferase bound to reconstituted discoidal ApoA1 HDL (11). However, little is known about the structural basis for the association of the many other protein components with HDL. Here, we present a structural model for lipid-bound and HDL-associated vitronectin (Vn), a multifunctional serum glycoprotein and component of the HDL proteome, found in the fraction of small, dense pre-β HDL particles associated with its acute phase response activities (5-8).

Vn interacts with a broad range of ligands to regulate cell adhesion and migration, extracellular matrix deposition, immune response, angiogenesis, fibrinolysis and more (12-15). It has been reported to bind lipids and cholesterol (16) and is found in the lipid-rich deposits that accumulate with diseases of aging, including atherosclerosis (17), age-related macular degeneration and Alzheimer’s disease (18). Understanding its mechanism for lipid binding is important for gaining insights to these physiological and pathological processes.

Discoidal HDL assemblies consist of two copies of the major structural protein ApoA1 bundled as an antiparallel helical double-belt around the hydrophobic edge of the lipid bilayer disc (19-23), but recent work (22, 24) has shown that while the core helices of ApoA1 form a stable bundle, ApoA1 is conformationally flexible, especially when bound to small, 8-10 nm diameter, discoidal HDL. The ApoA1 N- and C-termini are significantly more flexible, and cross-linking mass spectrometry combined with long time scale MD simulations (22, 24) show that the C-terminus of one or both molecules can flip back on itself creating fenestrations that expose the lipid acyl chains and can result in local distortions of the HDL lipoparticle. These long time-scale simulations provide new structural models that offer a platform for exploring the interactions of nascent HDL with the various proteins in its proteome.

In this study, we used nuclear magnetic resonance (NMR) experiments and molecular dynamics (MD) simulations to first identify the primary binding site for phosphatidylcholine (PC) in the hemopexin-like (HX) domain of Vn, and then develop models of VnHX associated with either individual PC molecules or a discoidal HDL lipoparticle. PC is one of the most abundant phospholipids of the HDL lipidome (8, 9). We find that an extensive surface-exposed hydrophobic groove in the VnHX domain is capable of binding both the short-chain phospholipid 07:0 PC (diC7PC) and the natural mixed-chain phospholipid 16:0-18:1 PC (POPC). Lipid binding is guided by interactions of the lipid polar headgroups and acyl chains with positively charged and hydrophobic residues that organize the bound lipids in a tail-to-tail bilayer-like arrangement within the VnHX groove. Additional MD simulations of the resulting structural model of POPC-bound VnHX, combined with the published MD structural model of discoidal HDL that exhibits fenestration (22), show that VnHX associates with HDL at the fenestration and in a lipid-mediated manner.

These findings suggest a mechanism for the association of Vn with HDL and provide insight to its accumulation with the lipid-rich deposits that form in age-related pathologies. They expand the range of protein-lipid interactions observed within the structurally and functionally diverse collection of HDL particles.

## RESULTS

### Lipids bind a long groove on the surface of Vn

Vn circulates as a 75 kDa N-glycosylated polypeptide. Its C-terminal HX domain, comprising ∼70% of the mature protein sequence, harbors binding sites for various ligands and folds as a four-bladed β propeller (**Fig. 1A; Fig. S1**) with each blade formed by one of four HX homology sequences (HX1-HX4) (25). The propeller channel occludes NaCl, and four Asp residues at the channel rim form a Ca^2+^ binding site (26, 27). The HX1-HX4 interface forms an elongated, surface-exposed, hydrophobic groove that is an attractive candidate for lipid binding (**Fig. 1B, C**). To estimate the lipid binding potential of VnHX we performed NMR experiments with diC7PC, a soluble short-chain mimetic of native phosphatidylcholine (PC). Our rationale for using short-chain diC7PC in these experiments was dictated by the water insolubility of physiological long-chain lipids, which in turn limits our ability to generate samples for solution NMR and other structural methods. Future experiments with long-chain lipid and solid-state NMR have the potential of providing additional useful information.

**Figure 1.**
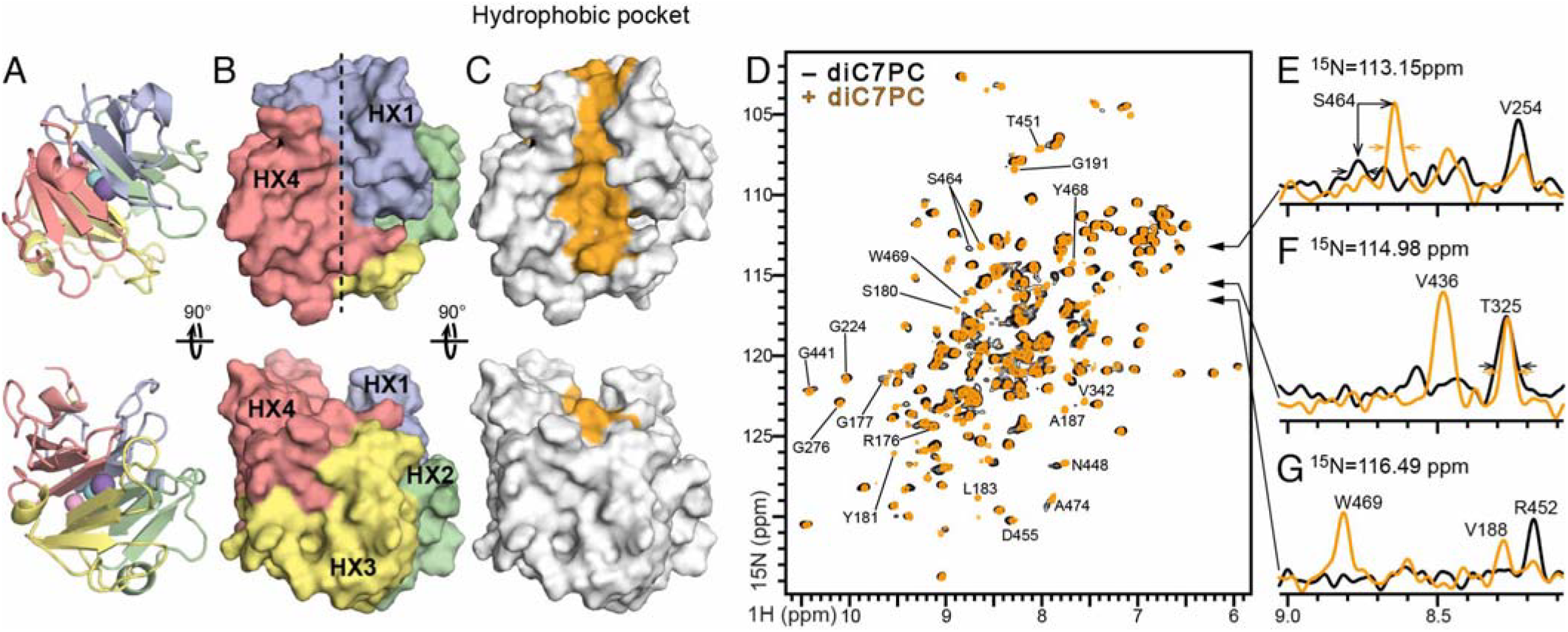
VnHX binds PC. **(A-C)** Ribbon and surface representations of VnHX. Throughout the study, VnHx colors denote repeats HX1 (blue), HX2 (green), HX3 (yellow) and HX4 (red), and the hydrophobic groove (dashed line; orange). Spheres denote Ca^2+^ (pink), Cl^−^ (aqua) or Na^+^ (indigo). (**D-G)** NMR 2D ^1^H/^15^N HSQC spectra (D) and 1D ^1^H spectra taken from the 2D (E), of ^15^N labeled VnHX acquired before (black) or after (yellow) addition of diC7PC (VnHX/diC7PC = 1/50 molar). Examples of new peaks (D, F, G), chemical shift perturbations (D, E), and line narrowing (F) observed upon addition of diC7PC are marked.

In previous NMR studies (26, 27) we showed that VnHX adopts the same β propeller structure in solution as in the crystalline state. Here, we find that diC7PC does not cause a major change of the ^1^H/^15^N NMR spectrum leading us to conclude that it maintains its overall structure. Addition of diC7PC, however, induces distinct ^1^H/^15^N perturbations, including changes in chemical shift, appreciable line narrowing and the appearance of many (∼37) new, previously undetected ^1^H/^15^N signals, representing nearly 20% of the total assignable backbone sites (**Fig. 1D-G; Fig. S2**). The NMR perturbations are highly specific, with the largest effects mapping to the surface-exposed hydrophobic groove of the HX domain and other regions of the protein structure remaining unaffected (**Fig. 2A-C**). The NMR line narrowing effects and appearance of new peaks are consistent with an overall increase of conformational order, indicating that lipid association stabilizes a specific state of the protein conformational ensemble. Overall, the NMR data show that lipids bind to the surface-exposed hydrophobic groove of the HX domain.

**Figure 2.**
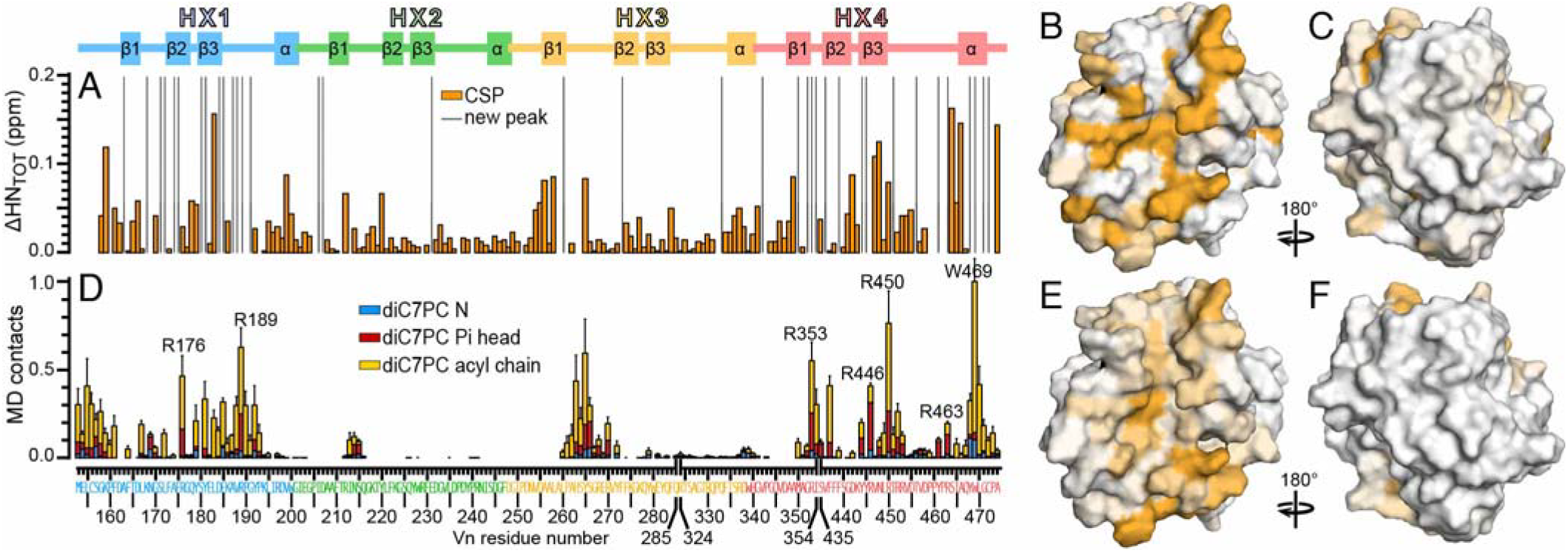
NMR and MD simulations map the lipid binding site of Vn. **(A)** NMR backbone amide ^1^H and ^15^N chemical shift perturbation (ΔHN_TOT_ = ΔH + ΔN/5) induced by diC7PC. The amino acid sequence (bottom) and protein secondary structure (top) are colored by each HX motif as in the structures. **(B, C)** Three-dimensional surface of VnHX colored to reflect low (white) to high (yellow) level of lipid-induced NMR perturbations. (**D)** Protein-lipid contact frequency from MD simulations. Data are averaged over all replicas. The bars represent the frequency of finding a heavy atom of each protein residue within 4Å of heavy atoms in the lipid acyl chain (yellow), or polar headgroup phosphate (red) and trimethylammonium nitrogen (blue). Error bars denote standard error of the mean. **(E, F)** VnHX surface is colored to reflect low (white) to high (orange) protein-lipid contact frequency derived from MD simulations.

The lipid binding affinity of VnHX may be estimated by monitoring the changes in ^1^H/^15^N chemical shifts as a function of increasing diC7PC concentration. The experimental data for well-resolved ^1^H/^15^N signals fit a lipid binding model (28) with an average apparent equilibrium dissociation constant (K_d_) of 68±25 µM and a binding stoichiometry of 4 lipids per protein (**Fig. S3**). In the native state, Vn is expected to associate with long-chain lipids which may exhibit higher affinity for the hydrophobic groove. Moreover, affinity estimates with diC7PC are complicated by its micelle forming ability, thus additional experiments with a broader range of lipid types and concentrations will be needed to accurately assess the lipid binding affinity. These limitations notwithstanding, the current data provide an estimate of lipid binding by the VnHX domain.

To gain deeper structural insights, we performed five independent 6 µs MD simulations for a total aggregate time of 30 µs, starting from the crystallographic coordinates of VnHX (PDB 7rj9) and 20 molecules of diC7PC dispersed in water (**Table S1**). Because the NMR experiments were carried out with diC7PC we sought to perform analogous MD simulations that would complement and cross validate the experimental data.In line with the NMR observation that diC7PC does not induce a major structural rearrangement of the VnHX β-propeller, the protein coordinates equilibrate within a few nanoseconds and the structure remains folded over the course of simulation (**Fig. S4**).

All simulations resulted in both rapid association of individual lipid molecules with the protein, and micellar self aggregation of the remaining unbound lipids (**Fig. S5**). Volumetric map analysis (**Fig. S6**) reveals the formation of a spherical ∼40 Å diameter micellar structure, with an average aggregation number of 16 lipids, that can be visualized as an inner volume of lipid acyl chains enclosed by an outer shell of polar headgroups, as expected for diC7PC in water. The analysis also identifies two types of lipid-protein interactions: specific binding of individual lipid molecules to the HX1-HX4 hydrophobic groove, and a more dynamic association of the micelle with Arg-rich surface sites near the ends of the groove and at the β1-β2 loop of HX3.

Notably, the pattern of protein-lipid contacts calculated for all heavy atoms from the MD trajectories (**Fig. 2D, E; Fig. S7A**) closely mirrors that of the experimental NMR chemical shift perturbations in identifying the HX1-HX4 groove as the primary lipid binding site. The excellent agreement of the MD simulations with the NMR data provides high confidence that the resulting MD structures are consistent with the experimental system. Moreover, while the MD atomic contacts reflect all heavy atoms in the lipid and protein, the ^1^H and ^15^N NMR signals report on changes in the chemical environment of protein HN backbone sites, induced by either direct lipid contacts with the backbone, or perturbations transmitted allosterically by lipid contacts with sidechains and more distal sites. In this way, the NMR and MD data are highly complementary, and provide a comprehensive view of the protein-lipid interactions.

The MD contact map also shows that Arg residues situated at each end of the groove (R176, R189, R353, R446, R450 and R463) make frequent contacts with lipid headgroup phosphates, while aromatic and hydrophobic residues within the groove contact the lipid acyl chains. Similarly, the NMR data reveal lipid-induced ^1^H/^15^N perturbations for the HN backbone sites of R176 and R450, and the appearance of new ^1^H/^15^N peaks for R189, R353 and R463 (no backbone assignments could be made for R446). Significant MD atomic contacts and NMR ^1^H/^15^N perturbations are also observed for aromatic residues that line the length of the hydrophobic groove forming an aromatic cage (F161, F164, Y181, F437, F439, Y444, and W469).

### MD simulations identify two lipid binding sites in the hydrophobic groove

In all MD simulations, individual lipids bind very rapidly, within the first 200 ns of simulation, as evidenced by a sharp reduction in the lipid-to-groove distances, and the groove remains lipid-bound for the duration of MD (**Fig. 3A; Fig. S5**). In all five replicas, initial lipid binding occurs at the HX4 end of the groove (designated site 1), guided by interactions of the lipid acyl chains with aromatic and aliphatic residues in the groove core, and polar interactions of the lipid phosphate group with three Arg (R353, R446, R463) that form a positively charged gate at the groove entrance (**Fig. 3A, C, pink**). This first lipid binding event is rapidly followed by a second lipid binding the HX1 end of the groove (designated site 2), guided by interactions of the lipid acyl chains with aromatic and aliphatic residues in the groove core, and polar interactions of the lipid phosphate group with R176 at the HX1 groove entrance (**Fig. 3A, D, blue**). Site 1 lipid binding always precedes site 2, but whether this sequence of events reflects cooperativity, with one event priming the other, cannot be discerned from the existing data.

**Figure 3.**
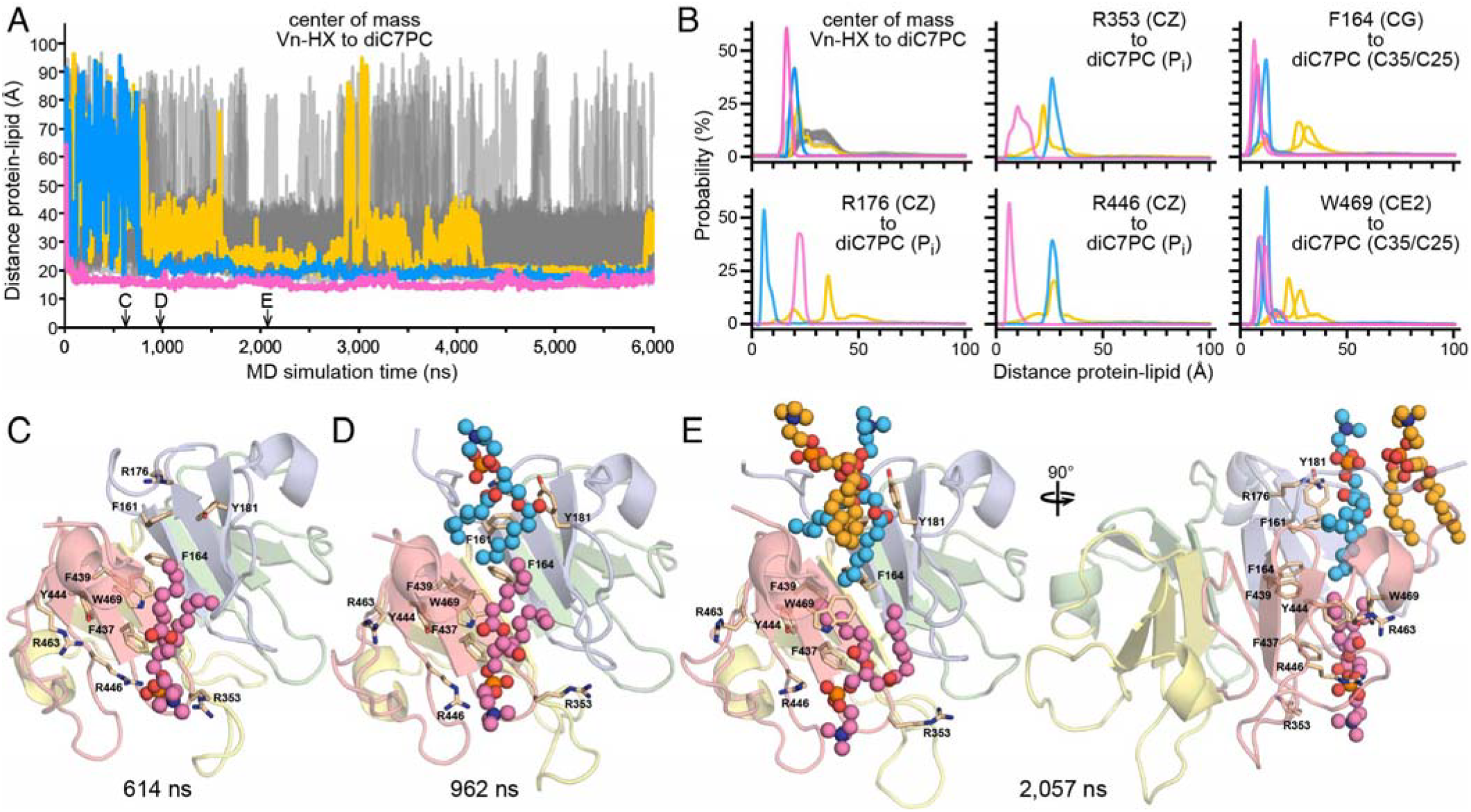
Short-chain lipid binding. **(A, B)** Representative time evolution (A) and probability distribution (B) of the center of mass and atomic distances between VnHX and each diC7PC molecule. Each trace represents a protein-bound (pink, blue and yellow) or micelle-bound (gray) lipid molecule. Arrows mark time points where snapshots (C-E) of the structure were taken. (**C-E)** Representative structures, taken at three different time points of MD simulation. Spheres (pink, blue, yellow) denote lipid bound at site 1 (pink), site 2 (blue) or recruited (yellow) from the pool of diC7PC. Sticks denote key aromatic and Arg sidechains.

Lipid binding at these primary sites is long-lived on the 6 µs time scale of MD. The two lipids occupy the groove in a tail-to-tail, bilayer-like conformation, with polar heads stabilized by phosphate-Arg contacts and acyl chains stabilized by the aromatic cage (F161, F164, Y181, F437, F439, Y444, W469) in the groove (**Fig. 3D**). Additional lipid molecules also associate with the groove-bound lipids in a way that is highly reminiscent of cooperative lipid bilayer formation (**Fig. 3A, E, yellow**). The site 1 and site 2 groove-bound lipids (**Fig. 3A, E, pink, blue**) are appreciably more conformationally restrained than either the additional surface-bound lipids or the micellar lipids, as evidenced by the narrow distribution of lipid-protein inter-atomic distances that they maintain over the course of simulation (**Fig. 3B**). By contrast, the surface-bound lipids (**Fig. 3, yellow**) exchange more frequently with the micelle and exhibit broad distributions of lipid-protein distances over the course of MD.

### Structural organization of the groove-bound lipids

To identify the dominant lipid-binding poses at the two primary groove sites, we analyzed a non-redundant set of structural descriptors that capture key conformational rearrangements of both lipid and protein upon binding (**Fig. 4**). Specifically, the minimum distances between lipid tail atoms and protein groove atoms from site 1 (F164 and M350) or site 2 (F161 and F164), each capture lipid binding at the respective sites. The time evolution of this minimum lipid-groove distance reflects a stable lipid binding interaction across all five replicas (**Fig. S8A**). The most probable occupancy of the groove corresponds to two bound lipids, with an average of 1.85 ± 0.5, with non-negligible probability of binding additional diC7PC molecules (**Fig. S8B**).

**Figure 4.**
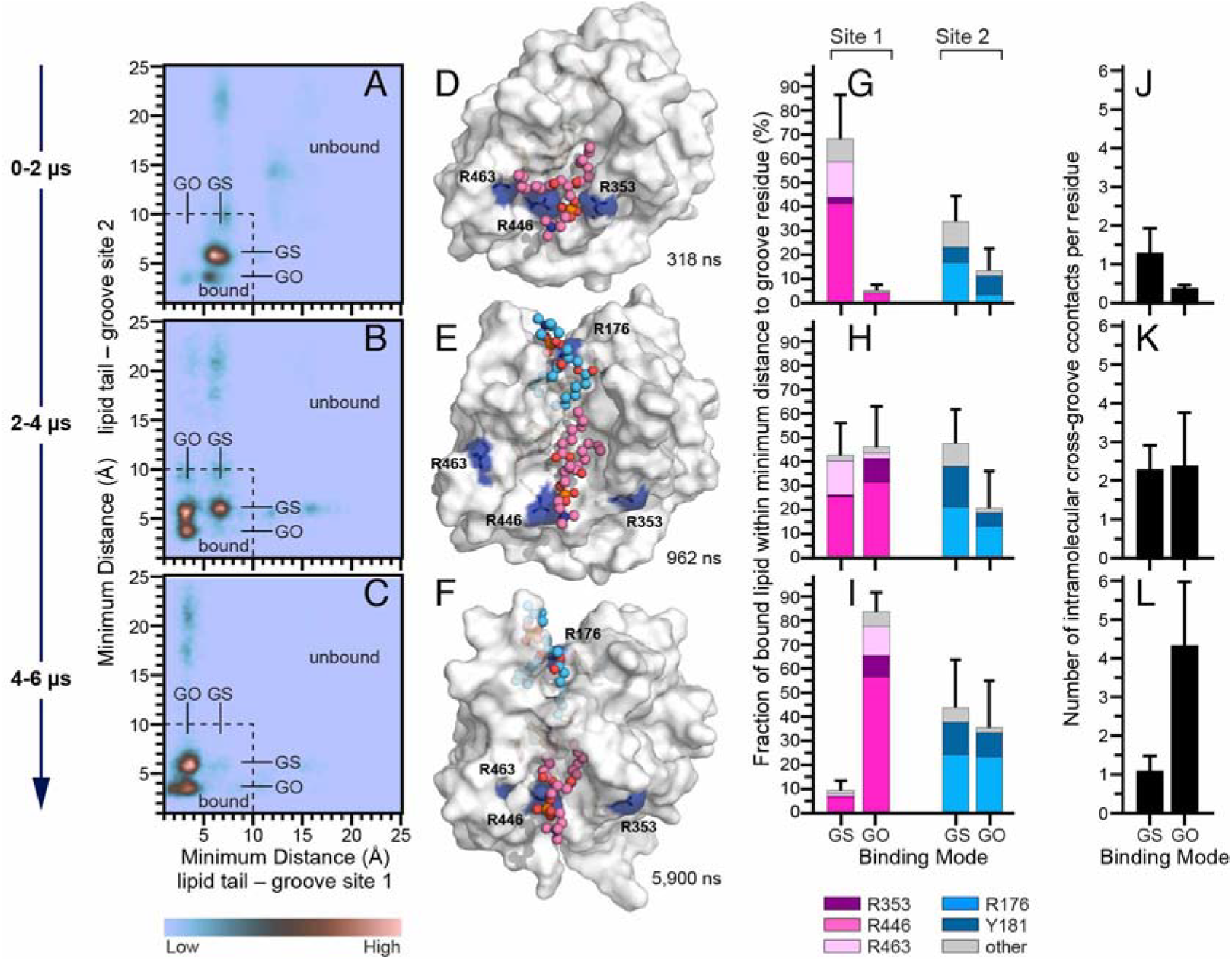
Cluster analysis of lipids bound to VnHX. The analysis was averaged over all replicas and performed at three different time blocks of simulation, from 0-2µs (A, D, G, J), 2-4µs (B, E, H, K), or 4-6µs, (C, F, I, L). Lipids are bound to the groove surface (GS), or occluded within its aromatic cage (GO). (**A-C)** Heatmap of probability density as a function of the minimum lipid distance to groove sites 1 and 2 (see Experimental Section). Blue to red color intensity reflects low to high lipid site occupancy. Site 1 lipids are clustered based on the minimum distance between lipid tail carbon atoms and the center of mass of the F164 CZ and M350 CE atoms. Site 2 lipids are clustered based on the minimum distance between lipid tail carbon atoms and the center of mass of the F164 and F161 CZ atoms. (**D-F)** Structural models representative of the GS and GO lipid binding poses. Blue surface denotes key gate Arg. Spheres (pink, blue, yellow) denote lipid bound at site 1 (pink) or site 2 (blue). (**G-I)** GS- and GO-bound lipid conformations exhibit distinct contacts with specific groove residues. GS and GO fractions represent the percentages of frames in each binding mode, with the remaining frames corresponding to unbound or loosely bound lipids; the three fractions sum to 100%. Within each binding mode, every frame is assigned to one of the indicated residues based on the shortest distance between its side chain and the lipid phosphate group at the respective site, provided the minimum distance is below 8 Å; otherwise, it is assigned to “Other.” **(J–L)** Clustered protein conformations exhibit distinct numbers of intramolecular atomic contacts across the groove, defined using a 7 Å cutoff between heavy atoms, that correlate with GS and GO lipid-binding poses. Error bars in panels G–I and J–L represent the standard errors of the total GS and GO fractions and the number of intramolecular contacts, respectively, calculated by dividing the frames into five equal blocks.

Mapped as a function of these minimum distance descriptors, the conformational landscape (**Fig. 4A-C**) reveals distinct peaks of population density, each corresponding to a groove-bound pose with the lipid acyl chains resting upon the groove’s aromatic cage (**Fig. 3D, 4E**), or a groove-occluded pose with one or both acyl chains inserted deep in the groove under the rings of the aromatic cage (**Fig. 3E, 4F**). The most probable states are those where both sites 1 and site 2 are occupied (**Fig. 4A-C**). For Site 1, bound and occluded poses form distinct peaks of the landscape separated by a low-probability transition region, reflecting long, µs scale residence times in each state with rare transitions between them. These states are correctly distinguished as separate structural groups by density-based clustering. For site 2, the separation between bound and occluded modes is less distinct than for site 1, with transitions occurring on the sub-µs scale.

Clustering the MD data in 2-µs time blocks further reveals the evolution of bound lipid populations over time. This analysis shows that the probability of simultaneously binding lipids at both sites 1 and 2, increases significantly in the last 2 µs. Initial lipid binding (0-2 µs) occurs predominantly in the groove surface bound mode, and gradually transitions to the occluded mode as the simulation progresses (2-4 µs). In the final stage (4-6 µs), the occluded mode dominates, suggesting that it drives the binding affinity, with the bound mode likely serving as a kinetic intermediate. Each pose correlates strongly with contacts of the lipid phosphate to specific residues in the Arg gates for either site 1 or 2 (**Fig. 4D-I**).

Viewed through the lens of the NMR data, the observations of two long-lived groove-bound lipids and the ability of additional lipids to associate more loosely, suggest that the Kd estimates from experimental NMR data reflect a mix of higher and lower affinity interactions.

### Conformation of lipid-bound VnHX

The NMR line narrowing effects of diC7PC, including the appearance of new peaks, indicate that lipid binding stabilizes a specific conformational state of the protein. In MD simulations, the transition from groove-surface to groove-occluded poses is associated with rearrangements of the W469 aromatic ring at site 1 and the Y181 aromatic at site 2 (**Fig. 3C-E**) and, for both these sites, diC7PC induces significant ^1^H/^15^N perturbations in NMR and establishes extensive atomic contacts in MD simulations (**Fig. 2A, D**). To explore the nature of such lipid-induced conformational bias, we monitored intramolecular protein-protein contacts between two groups of residues, each located on opposite sides of the hydrophobic groove: residues 155, 469–470 and residues 186–192 (**Fig. 4J-K**). Averaged over all replicas, across 2 µs time blocks, these cross-groove contacts increase in number as the protein transitions from the groove-surface to the groove-occluded state that dominates in the final 2 µs.

Finally, the MD simulations and NMR data combined indicate that these subtle conformational changes induced by lipid binding are transmitted beyond the lipid binding groove to distal sites. For example, new NMR peaks or significant chemical shift perturbations, but no MD atomic contacts, are observed for residues in blades HX2 and HX3, located polar opposite to the groove (I206, D207, E231, V254, A256, L258, F273, F284, V342), suggesting that the change in chemical environment sensed by NMR is a result of conformational change rather than direct interaction with lipid **(Fig. S9)**.

### Association with native long-chain lipids

Polyunsaturated fatty acids, including 18:1 carbon chains, are found to be enriched in the pre-beta-HDL particles (8) where Vn is also enriched. To explore how Vn might bind such long-chain native lipids, we initiated a new set of MD simulations in which Vn-bound diC7PC molecules were replaced by the mixed chain phospholipid POPC. The lipid molecules share the same PC headgroup and glycerol backbone, and differ only in acyl chain length and saturation. We considered three possible binding configurations: a single POPC molecule occupying either groove site 1 (VnHX–POPC_s1_) or site 2 (VnHX–POPC_s2_), and two POPC molecules one in site 1 and the other in site 2 (VnHX–POPC_s1s2_). These three starting structures (**Table S1**) were generated by gradual alchemical transformation (29) of Vn-bound diC7PC into POPC, and then used to initiate two independent 4.5-µs MD simulations for each state, for a total aggregate MD simulation time 18 µs.

In all simulations, POPC remains bound to VnHX for the duration of MD (**Fig. 5A-C**) and the map of atomic contacts identifies the VnHX groove as the binding site (**Fig. S7**). As observed for diC7PC, the lipid phosphates make polar contacts with the Arg gates at each end of the groove, and the acyl chains are stabilized by aromatic and aliphatic residues in the groove core. POPC initially bound to site 1 stays bound to site 1 (**Fig. 5A, D**). By contrast, POPC initially bound to site 2 transfers to site 1 (**Fig. 5B, E**), confirming the site 1 preference observed for diC7PC. Notably, two POPC molecules, each initially bound to site 1 or 2, also remain bound (**Fig. 5C, F**). As for diC7PC, they adopt a tail-to-tail conformation, although the longer acyl chains of POPC are intercalated adapting to the ∼25 Å length of the groove which is set by the distance between the Arg gates at sites 1 and 2. The data add to the physical evidence for native lipid association with the VnHX groove.

**Figure 5.**
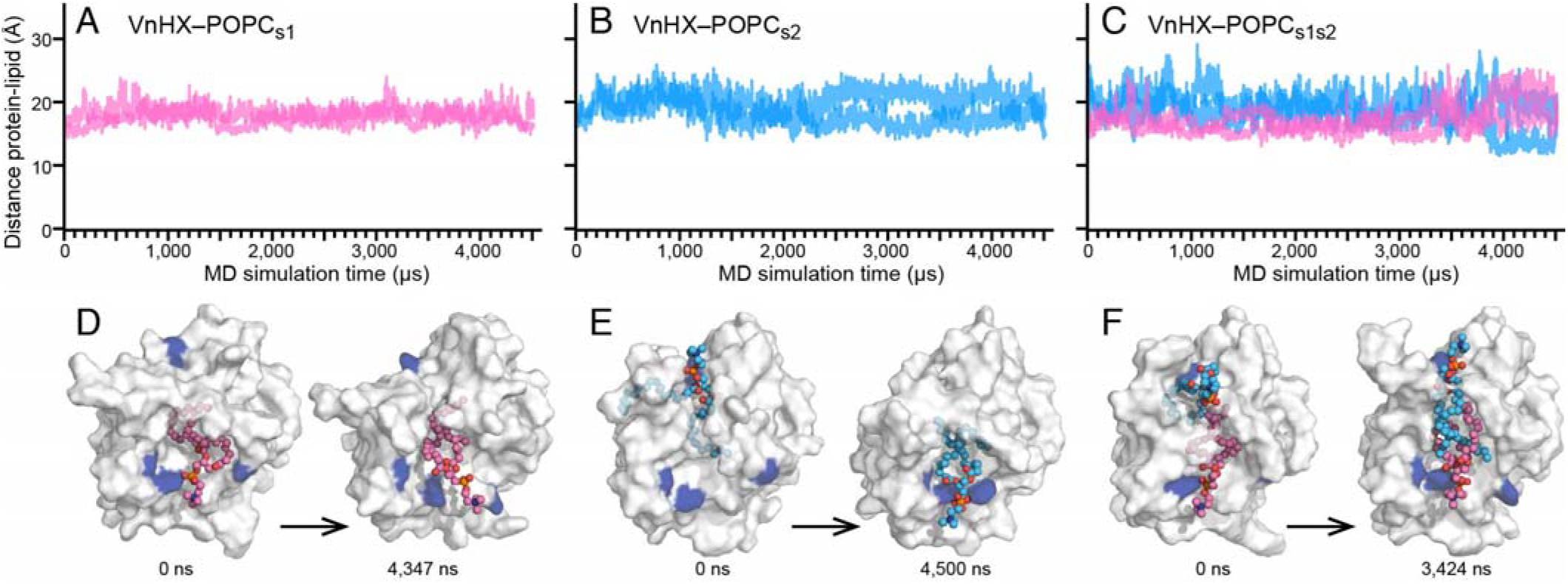
Long-chain lipid binding. **(A-C)** Representative time evolution of the distance between the centers of mass of VnHX and each POPC lipid molecule. Each trace was obtained for a starting model with one POPC bound to site 1 (A, pink), one POPC bound to site 2 (B, blue), or two POPC molecules each bound to site 1 and site 2 (C). (**D-F)** Representative structures, taken at the first (0 µs) and advanced time points of 4.5 µs MD simulation. Spheres (pink, blue, yellow) denote lipid bound at site 1 (pink) or site 2 (blue). Blue surface denotes key gate Arg.

### Association with an HDL lipoparticle

The long time scale MD models of HDL show that ApoA1 is flexible and its C-terminus can flip away from the regular double-belt to expose the lipid bilayer edge of the discoidal particle (22, 24). Could the association of Vn involve the exposed edge of such a partially de-scaffolded particle? To explore this question, we performed MD simulations of POPC-bound VnHX with a discoidal HDL lipoparticle containing POPC and ApoA1 (160:2 stoichiometry). Two starting models were generated by docking VnHX-POPC_s1_ to one of two HDL structures selected from the published MD trajectories (22): One, taken at 20 µs of the trajectory, is more canonically discoidal, whereas the other, taken at 10 µs, is less distinctly discoidal and more distorted. Both starting structures exhibit fenestration (**Fig. S10**). Together, they introduce HDL conformational variability and provide distinct environments for assessing the association of VnHX. Each starting model was equilibrated and subjected to three independent 5 µs simulations for a total aggregate MD simulation time of 30 µs.

VnHX remains associated with the HDL lipoparticle in all simulations (**Fig. 6A, B**) but one (**Fig. 6C**) where it exchanges between associated and disassociated states. The primary binding site is at the HDL fenestration where the C-terminal helix of one ApoA1 flips back to expose the lipid bilayer (**Fig. 7**). In all six simulations, including upon HDL dissociation, VnHX retains a POPC molecule bound in its groove. The map of atomic contacts (**Fig. S7E**) mirrors the data obtained for diC7PC and POPC and identifies the groove as the lipid binding site in all three models (diC7PC, POPC, HDL). The same POPC molecule that is bound to groove site 1 in the starting model (POPC_s1_) remains bound in all but one replica (**Fig. 6B**) and additional POPC molecules from the HDL pool also associate with the original POPC in the VnHX groove (**Fig. 6D, E**). Interestingly, the lost POPC_s1_ incorporates in the HDL pool and is in turn replaced in the VnHX groove by another POPC from the HDL pool (**Fig. 6F**). The data thus show that POPC can exchange between VnHX and HDL.

**Figure 6.**
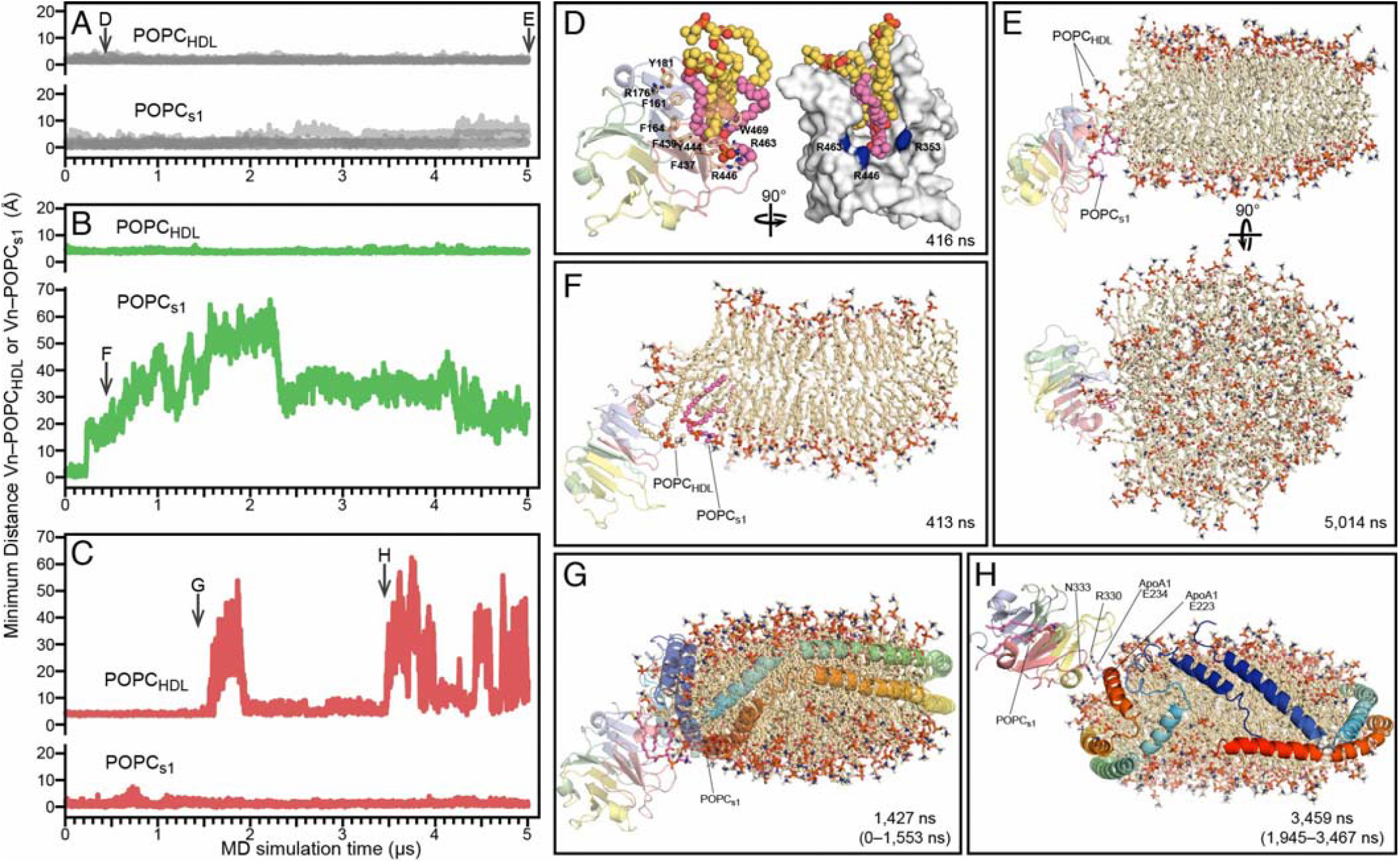
VnHX–HDL association. **(A-C)** Time evolution of the minimum distance between the center of mass of VnHX to the the center of mass of HDL POPC (POPC_HDL_) or of its bound POPC molecule (POPC_s1_). Arrows mark time points where snapshots (D-H) were taken for each trace. **(D)** Structure of VnHX taken from (A) showing bound lipids in the groove. Key residues in the aromatic cage and the Arg gates are marked. **(E)** VnHX bound to the HDL fenestration via its original POPC^s1^ and two additional POPC molecules from the HDL pool. **(F)** Exchange of the original POPC_s1_ with the HDL lipid pool. **(G, H)** VnHX spends ∼1.5 µs bound to the HDL fenestration (G) and ∼1.5 µs loosely associated with the C-terminal helix of ApoA1 via polar contacts (H), before dissociating from HDL. ApoA1 is removed from the image (E, F) or shown as cartoon (G, H). ApoA1 blue-red color gradient denotes N to C terminus progression. POPC (sphere) color denotes the original POPC_s1_ (pink), or POPC_HDL_ pool (wheat).

**Figure 7.**
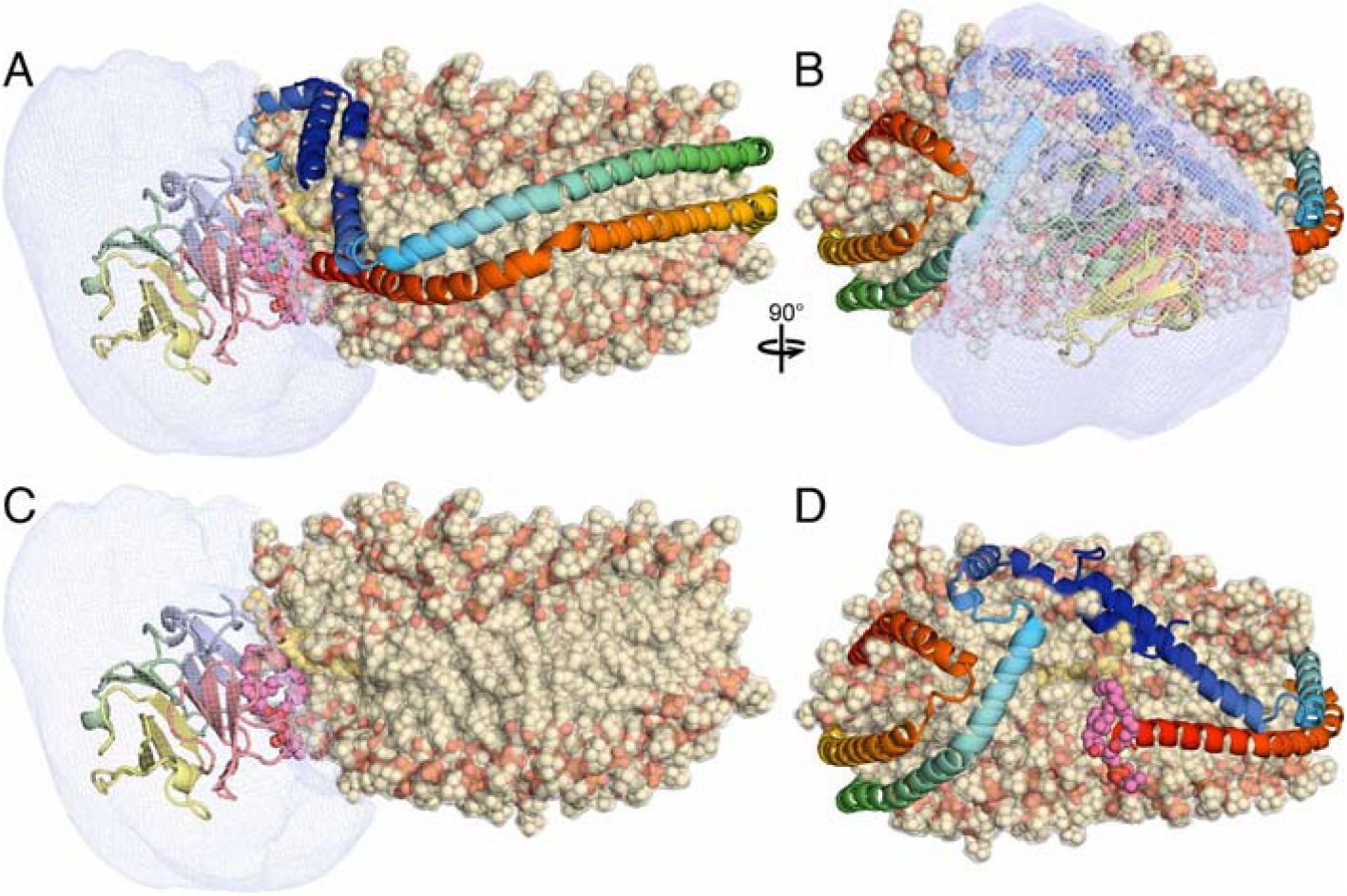
VnHX–HDL association, average model. **(A, B)** Orthogonal views of the average VnHX–HDL structure with volume density (mesh) of the time-averaged position of VnHX across all six replicas. **(C, D)** Same view as (A, B) with either ApoA1 removed to show the VnHX and HDL lipids (C), or VnHX removed to show the fenestration (D). VnHX (cartoon) color as in other Figures. ApoA1 blue-red color gradient denotes N to C terminus progression. POPC (sphere) color denotes the original VnHX-bound molecule (pink), or POPC of the HDL pool (wheat).

An additional mode of HDL association is observed in one of the six replicas (**Fig. 6C, G, H**). Here VnHX transiently associates with the flipped-back C-terminal helix of ApoA1, stabilized by polar contacts VnHX R330 with ApoA1 N333/E234, and VnHX R452 with ApoA1 E223, but these interactions are more transient and do not result in stable reassociation. This further supports the ability of the VnHX groove to stably bind lipids independently of the HDL particle. The transient nature of the alternative protein-protein interactions, together with the stable lipid-binding capacity of the groove, and the sustained association with the HDL fenestration, all point to a substantial contribution from lipid-mediated interactions.

The time-averaged volumetric map calculated over all six replicas identifies the HDL lipoparticle fenestration as the primary binding site for VnHX (**Fig. 7A**). As reported previously (22, 24), marked fenestration is produced when the C terminus of one ApoA1, and the N termini of both ApoA1 molecules, flip away from the rim of the HDL disc, exposing the lipid acyl chains to the aqueous milieu. The combined MD data show that the interaction of VnHX with the HDL fenestration is dynamic and allows for some range of binding poses localized at the site. Pronounced local deformation of the HDL lipid bilayer at the fenestration is evidenced by bending and reorientation of the lipid headgroups toward the aqueous solvent. This likely provides a favorable environment for the interaction with VnHX, as protein binding could reduce or stabilize this local deformation by lowering the energetic cost associated with water exposure of the acyl chains, thereby contributing driving force for its association.

Overall, the persistence and dynamic nature of the association support a model in which VnHX associates with HDL via lipid-mediated interactions at the fenestration site. Additional stabilization may arise from VnHX-ApoA1 interactions, including observed polar contact sbetween VnHX R189 and ApoA1 D48 and D51. This electrostatic contribution to binding may be important for initially guiding VnHX towards the lipoparticle, but appears to play a minor role in maintaining VnHX bound to the fenestration, given its marked localization at this ApoA1-free site.

## DISCUSSION AND CONCLUSIONS

Recent studies indicate that the function of HDL transcends the traditional correlation of its cholesterol level with risk of atherosclerotic cardiovascular disease. While HDL is well known for its central role in removing excess cholesterol from cells and transporting it to the liver for excretion, it is also associated with other fundamental physiological processes, and HDL particles have diverse compositions, structures and sizes, tailored for specialized physiological functions (2-4). Proteomic studies have identified more than 280 HDL proteins that confer specialization, highlighting the need to understand the molecular basis for their association with HDL. Our work sheds light on the mechanism for HDL association by the HDL component glycoprotein Vn.

The NMR and MD data identify an elongated hydrophobic groove on the surface of the VnHX domain as the primary binding site for short-chain and long-chain phospholipid. Positively charged Arg gates at the ends of the groove, and an aromatic cage within the groove core, guide lipid binding in a tail-to-tail arrangement reminiscent of a lipid bilayer. Short-chain and long-chain PC molecules remain associated with the groove over extended periods on the microsecond time scale of MD simulations, and help engage additional PC molecules from the HDL lipid pool, as supported by the non-negligible probability of higher-order lipid occupancies within the Vn groove. The model of VnHX bound to HDL is consistent with the NMR data, as well as with previous experiments and MD simulations (22, 24) showing that HDL lipids are partially de-scaffolded at the fenestrations that form when the ApoA1 termini flip away from the classical double belt. This fenestration is the principal binding site for VnHX and may also offer binding opportunities for other HDL proteins with lipid binding capacity.

While the simulations do not permit determination of the absolute residence time, the absence of dissociation over the simulation trajectories indicates that the VnHX-HDL assembly persists beyond the microsecond time scale. The data point to preferential association with structurally perturbed regions of the lipoparticle, where local deformation and exposure of the lipid bilayer provide a favorable binding environment. The Arg gates of the groove set the phosphate P-P atomic distance of bound PC in the range of 25 Å, approximately 10 Å shorter than the ∼40 Å thickness of the HDL lipid bilayer, and VnHX binding correlates with edge remodeling of the HDL disc at the fenestration. In this way, Vn may help stabilize the deformed HDL regions through protein-lipid interactions, potentially promoting the association and stabilization of small or structurally perturbed HDL particles in the ∼90–100 Å size range. Alternatively, we cannot exclude the possibility that Vn binds only one or a few lipids circulating as the very high-density HDL that has been proposed (1). Chemical composition analysis of HDL-associated Vn and structure determination will be needed to answer this question.

HX domains are found in other protein families, most notably hemopexin (Hpx) and the matrix metalloproteases (Mmps). The latter have been reported to associate with membrane surfaces, including via their HX domains (30-32), and the zymogen of Mmp2 was recently reported to associate with ApoA1 through its HX and catalytic domains (33). In all these cases, the protein interactions are membrane-peripheral and individual lipid binding has not been reported. Hydrophobic residues at the HX1-HX4 interface are conserved across the sequence of Vn, Hpx and MMPs, but whether the HX1-HX4 interface functions as a lipid binding site in other proteins remains an open question.

Ultimately, lipid binding, including the proposed affinity for cholesterol, and the surface accessibility of the groove will need to be validated in the context of native full-length Vn and long-chain lipids. The AlphaFold structural model of full-length Vn predicts that the groove is accessible for lipid binding, unobstructed by other protein segments or N-glycosylation at N169 and N242 (**Fig. 8**). Whether additional regions of the protein contribute to lipid binding will have to be determined experimentally. Moreover, the VnHX groove is stabilized by a C156-C472 intramolecular disulfide bond, and disulfide bond reduction could alter the groove’s lipid affinity and specificity, but Vn is reported as a substrate of protein disulfide isomerase and disulfide bond reshuffling affects its interactions with integrin receptors (34), thus, redox-dependent conformational changes in Vn may also govern HDL association.

**Figure 8.**
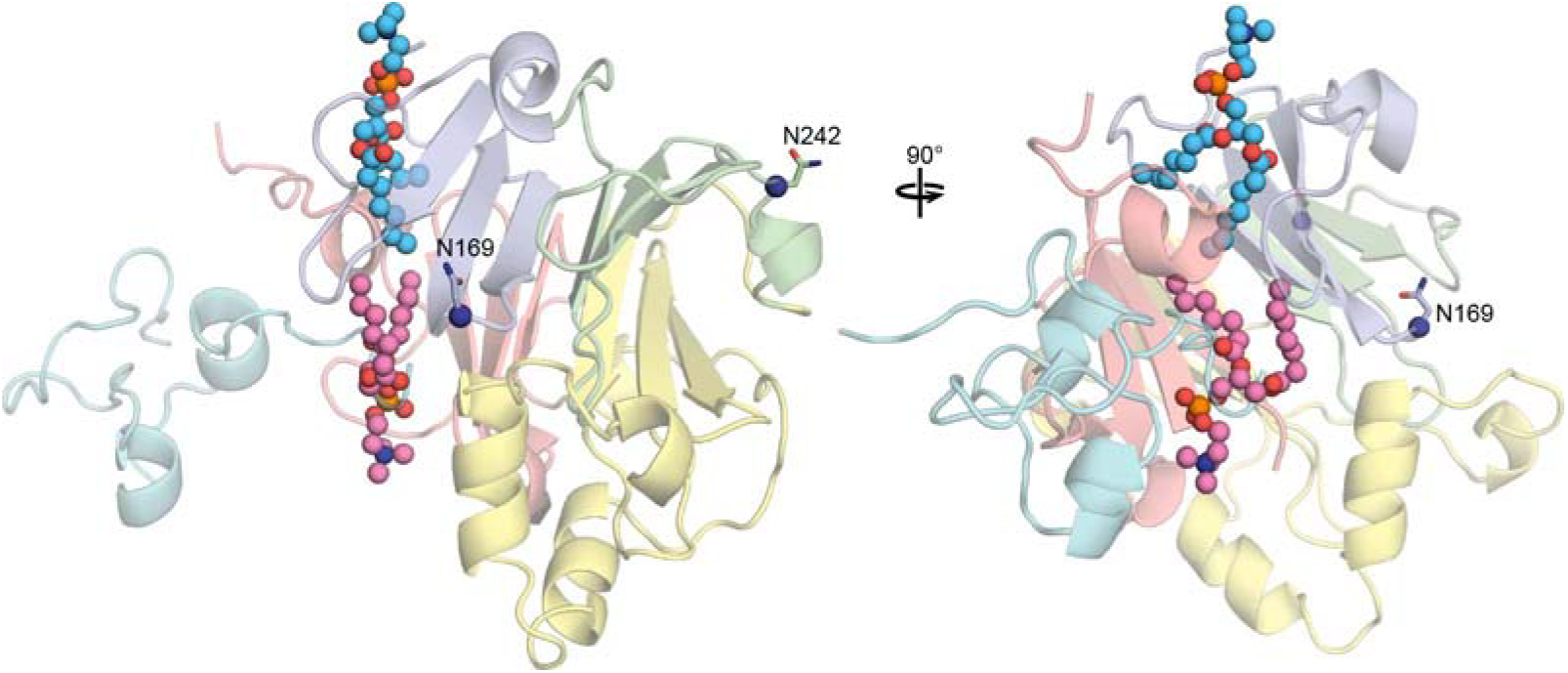
Model of lipid binding in the context of full-length Vn. The AlphaFold model of Vn was superimposed with a representative structure of VnHX taken at 2,057 ns of MD simulation, with bound lipid molecules (pink, blue spheres) bound to the HX1-HX4 cleft. Long unstructured regions not predicted by AlphaFold are not shown. Protein colors reflect: the SMB domain (cyan), HX repeats (HX1, blue; HX2, green; HX3, yellow; HX4, red), and the HX3 helical extension (yellow). Two N-glycosylation sites (N169, N242) are marked.

Finally, Vn is present at relatively high concentrations in human plasma. While the present data do not allow us to estimate the proportion of total plasma Vn carried by HDL, its apparent affinity for lipid may suggest that Vn circulates at least as a lipid-associated protein, and its association with lipid and HDL has physiological implications for human health.

Vn circulates in blood and is found in the extracellular matrix of various tissues, but it is also a major constituent of the subretinal, lipid-rich, calcified deposits that accumulate with age related macular degeneration (AMD) (35-38) and is found deposited in Alzheimer’s disease brains (18). Retinal pigment epithelial (RPE) cells from AMD human donors have elevated levels of Vn protein, Vn protein levels are upregulated in complement-stimulated, immortalized RPE cells (39), and Vn mRNA is elevated in human induced pluripotent stem cells RPE cells derived from AMD patients (35-38).

In all these settings Vn is found associated with lipid and AMD is widely known to correlate with altered lipid metabolism across the visual cell layers (40). Neutral lipids represent a major component of age-related deposition in the eye and multiple biochemical, histochemical, and ultrastructural methods indicate that RPE cells secrete ApoB lipoparticles in the Bruch’s membrane – the sub-retinal extracellular matrix supporting the RPE layer. Vn-rich lipoparticles accumulate at this site, ultimately forming a precursor of the basal linear deposit hallmark of AMD. Components of these lesions have been proposed to interact with reactive oxygen species to form pro-inflammatory peroxidized lipids that induce neovascularisation.

The affinity of Vn for Ca^2+^, as well as lipid, is also relevant in the context of atherosclerosis and other pathologies characterized by calcified protein-lipid deposition. Phospholipids have inherent affinity for Ca^2+^ ions and can nucleate calcium-phosphate clusters to initiate mineralization (41). Thus, by simultaneously recruiting both lipids and Ca^2+^, Vn may play a role in mediating calcium-phosphate mineralization, as proposed previously (26).

Vn is at the intersection of genes that associate with AMD and are also historically associated with plasma HDL. The model for Vn-lipid and Vn-HDL association is a provides a framework for exploring the role of Vn in HDL function and models of HDL protein association and a basis for future research directions and therapeutic strategies based on its lipid-binding property.

## EXPERIMENTAL METHODS

### Protein preparation

VnHX was prepared from *E. coli* as described previously (25). Uniformly ^15^N/^13^C labeled protein for NMR studies was obtained by growing bacteria on minimal M9 media containing (^15^NH_4_)_2_SO_4_ and ^13^C-glucose (Cambridge Isotopes). Samples for NMR contained 40-164 µM VnHX in 160 μL of buffer (20 mM MES, pH 6.5, 300 mM NaCl, 2 mM CaCl_2_) supplemented with 5% D_2_O. Lipid containing samples were supplemented with 1,2-diheptanoyl-sn-glycero-3-phosphocholine (diC7PC) to obtain a protein/lipid ratios of 1/1, 1/10, 1/50 and 1/100.

### NMR Spectroscopy

Solution NMR experiments were performed at 30°C on a Bruker Avance 600 MHz spectrometer, equipped with a ^1^H/^13^C/^15^N triple-resonance cryoprobe. The NMR data were processed with TopSpin (Bruker) and analyzed using CCPN 2.4.2 (42). The HN, N, and CA chemical shifts were assigned using two-dimensional ^1^H/^15^N heteronuclear single quantum coherence (HSQC) experiments, and three-dimensional ^1^H/^15^N/^13^C HNCA experiments (43). The assigned chemical shifts are deposited in the BMRB database (IDs: 50241, 50261, 53399). Values of the apparent dissociation constant (Kd) were estimated by fitting the NMR data to an n-site binding model, as described (28), with the equation:

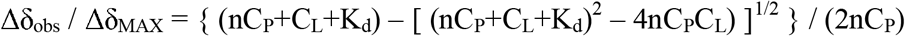

where Δδ_obs_ is the observed change in chemical shift, Δδ_MAX_ is the maximum change observed at saturation, n is the number of binding sites, C_P_ is the protein concentration (40 µM) and C_L_ is the lipid concentration (0-4 mM).

### Molecular dynamics (MD) simulations

All-atom MD simulations were performed as described (27), using the CHARMM36 force fields with the TIP3P water model (44). The temperature was set to 303.15K for simulations at 30°C, and the pressure was maintained at 1 bar. The systems for MD (**Table S1**) were prepared and equilibrated using the CHARMM-GUI *Membrane Builder (45)*. The initial model of VnHX was generated from the Ca^2+^ bound crystal structure (PDB code 7rj9, molecule A) (25), after removing all ligands and water molecules, and modeling residues with missing electron density using GalaxyFill (46). Statistical analyses were performed with home-made Python scripts in MDAnalysis (47), PyLipid and JupyterNotebook.

### Statistical Analysis

Trajectories were generated every 1 ns for 6 µs (DHPC), 4.5 µs (POPC), and 5 µs (LPP) and the last 5.8 µs (DHPC), 4.3 µs (POPC), and 4.8 µs (LPP) were used for statistical analysis with home-made Python scripts in MDAnalysis (47), PyLipid and JupyterNotebook.

For atomic contact analysis, intermolecular contacts for heavy atoms within 4 Å of each other were considered in each frame. Per-residue contact frequencies were computed independently for each replicate MD trajectory using a custom Python script built on MDAnalysis. Prior to analysis, trajectories were wrapped appropriately to ensure intact chains and correct treatment of periodic boundaries, then aligned to backbone of VnHX (DHPC and POPC data) and ApoA1 (LPP simulation). The first 200 frames of each simulation were omitted from analysis, corresponding to the first 200 ns. The resulting values were averaged across all replicas and the standard error of the mean calculated across replicates.

For conformational landscape analysis, to identify the most probable diC7PC binding poses within the Vn groove, we evaluated a set of structural descriptors capturing key electrostatic and hydrophobic protein–lipid contacts. The ability of these descriptors to distinguish distinct binding poses was assessed using the Density-Based Spatial Clustering of Applications with Noise (DBSCAN) algorithm (48-50), as implemented in the Scikit-Learn package (51), across a range of neighborhood distance and minimum sample parameters. Irrelevant or redundant descriptors were systematically filtered out, yielding two key non-redundant descriptors for lipid binding at sites 1 and 2, respectively. For site 1, this descriptor corresponds to the minimum distance between lipid tail carbon atoms and the center of mass of the F164 CZ and M350 CE atoms. For site 2, it corresponds to the minimum distance between lipid tail carbon atoms and the center of mass of the F164 and F161 CZ atoms. These two descriptors were then used to map the conformational landscape sampled by the MD simulations, clearly delineating the major binding states. Error bars (**Fig. 4**) were estimated using block analysis. For each selection, the total number of frames was divided into five blocks of equal size, and the standard error was calculated from the block averages.

### MD of Vn with diC7PC

The system for MD (**Table S1**), containing one protein and twenty diC7PC molecules, was used to initiate five independent MD production simulations, conducted with OpenMM using a time step interval of 4 fs with mass repartitioning. Trajectories were generated every 1 ns for 6 µs, and the last 5.8 µs were used for analysis.

### MD of Vn with POPC

Three systems (**Table S1**), each containing one protein and one or two POPC molecules, were generated for MD by means of an alchemical transformation (29) where Vn-bound diC7PC molecules were gradually transformed into POPC using the COLVARS library (52) and NAMD (53).

All three systems were generated from the same initial state, with two bound diC7PC molecules each in their respective s1 or s2 binding sites, selected from the conventional diC7PC simulations. One or two POPC molecules were added and aligned with the bound diC7PC via their common atoms. During the course of transformation, interactions between appearing (POPC) and disappearing (diC7PC) molecules were excluded, and non-bonded interactions of diC7PC and POPC were each gradually ramped down to zero (diC7PC) or ramped up (POPC). This process was controlled by the alchemical parameter λ, which was increased from 0 to 1.

To maintain spatial correspondence between the common atoms of POPC and diC7PC, pairwise distance restraints were applied with the harmonic potential :

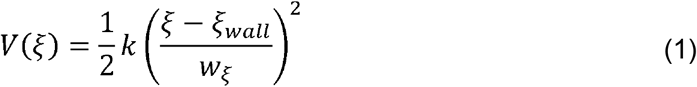

where *k* is the force constant, *ξ* is the instantaneous distance, _wall_ is the position of the harmonic wall, and is the characteristic width of the collective variable. To ensure precise overlap of corresponding C atoms, the harmonic restraint was centered at 0 Å, with a width of 0.05 Å, and a force constant of 5 kcal/mol. The system was first equilibrated for 11 ns at λ=0 to optimize the POPC–diC7PC overlap. The alchemical transformation was then performed over 50 sequential windows, over a total of 250 ns, with a λ ramp from 0 to 1. All diC7PC molecules were removed after the transformation, and the final transformed configuration was used to initiate two independent 4.5-µs MD simulations for each.

### MD of Vn with an HDL-like lipoparticle

Two conformations (**Table 1**) were selected from the MD trajectories reported previously (22) for a ∼100 Å recombinant HDL lipoparticle (rHDL-2-100) containing two molecules of the ApoA1 scaffold protein, 160 molecules of POPC, and 24 molecules of unesterified cholesterol (Zenodo: https://zenodo.org/records/897027).

Both systems exhibit discoidal lipoparticle structures with marked fenestration, produced when the termini of one or both ApoA1 molecules kink away from the lipid bilayer edge of the disc to expose the hydrophobic acyl chains. One system, selected at 10 µs of the trajectory, is a more distorted lipoparticle disc and more pronounced fenestration, while the second, selected at 20 µs, is a more regular disc (**Fig. S10**).

For each conformation, a model of Vn bound to one POPC molecule at site 1 (POPC_s1_) was selected from the MD trajectories of VnHX-POPC, and positioned near the lipoparticle fenestration with the POPC molecule aligned to the lipoparticle lipids. The resulting systems were solvated and neutralized using GROMACS (54) in a rhombic dodecahedral simulation box containing ∼0.5 million atoms. The MD simulations were performed in two steps, the first restrained and the second restraint-free. Both stages were performed at 298 K.

Restrained MD simulations were first performed to promote lipid-mediated association of Vn with the lipoparticle. In this step, the coordination between the Vn-bound POPC molecule and lipoparticle lipids was progressively increased during a restrained MD equilibration. Three collective variables (CV1-CV3) were used to control this process through the COLVARS library (52).

CV1 maintains the bound state of POPC within the Vn binding pocket. It corresponds to the coordination number between the bound POPC molecule and Vn, and was restrained to a value of 50, as estimated during equilibration.

CV2 progressively promotes the association between the Vn-bound POPC molecule and the lipoparticle lipids. It corresponds to the coordination number between Vn-bound POPC and the nearest lipoparticle lipids, defined as POPC or cholesterol molecules containing at least one heavy atom within 25 Å of Vn-bound POPC. The target coordination number was increased stepwise, over 100 ns, from 2 to 400 for the distorted lipoparticle system, and from 0.8 to 400 for the flattened system. The coordination number between two atom groups was calculated as:

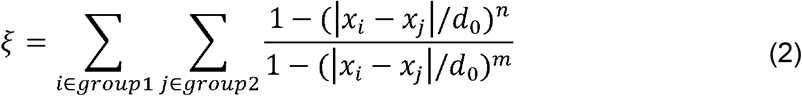

where x_i_ and x_j_ are the positions of atoms i and j, d_0_ is the cutoff distance, and n and m define the exponents of the distance-dependent switching function. Here, d_0_ was set to 5 Å, with n=6 and m=12.

CV3 maintains Vn in proximity of the lipoparticle during the restrained simulation. It corresponds to the distance between the centers of mass of Vn and the lipoparticle, and applies an upper-wall restraint ranging from 65 to 70 Å during the restrained simulations.

The three CVs were restrained with harmonic wall potentials (**Eq. 1**), with force constants of 0.1 kJ/mol. The wall widths were 0.1 for the coordination-number restraints and 5 Å for the distance restraint, allowing limited fluctuations while maintaining the desired association.

Equivalent harmonic wall potentials were applied to RMSD CVs calculated for the carbon atoms of Vn and lipoparticle, including both protein and lipid atoms, using their respective initial structures as references. The RMSD restraint widths were set to 1 Å (Eq. 1), with a harmonic center of 0.0 and a force constant of 1000 kJ/mol to preserve the structural integrity of both components during the restrained association simulations.

For the final step, the configuration obtained from restrained MD was used to initiate independent production restraint-free MD simulations. All restraints were removed, velocities were randomized, and the systems were simulated for 5 μs, with three independent replicas for each of the flat and distorted disc conformations.

Vn-lipoparticle association, and retention of the initial Vn-bound POPC molecule within the Vn binding pocket, were each monitored by measuring the distance between every fourth Vn CA atom and all C atoms of the lipoparticle lipids, or all C atoms of the initial Vn-bound POPC molecule, and the minimum distance reported over time.

## ASSOCIATED CONTENT

This article contains supplementary information (Table S1; Figures S1-S10).

## ACKNOWLEDGMENTS

We thank Marassi lab members for useful discussion, and Richard Pastor for sharing the long time scale MD trajectories of HDL. We also thank the reviewers for contributing useful insights. This research was supported by grants from the National Institutes of Health (P01 AG081167, R35 GM118186,

P30 CA030199), the Canadian Institutes of Health Research (201711MFE-395794-210656), and the Fishman Fund Postdoctoral Fellowship. Computing resources were supported by the Research Computing Center at MCW.

## AUTHOR CONTRIBUTIONS

KS, FM, FMM: writing, reviewing and editing KS: protein preparation, NMR experiments

KS, WB, YT, GT, FM, FMM: MD simulations and analysis

## Abbreviations

ApoA1: apolipoprotein A-I
diC7PC: diheptanoyl-phopshatidylcholine
HDL: high-density lipoprotein
HX: hemopexin
MD: molecular dynamics
NMR: nuclear magnetic resonance
Vn: vitronectin

