## supporting information for "Structural model of the blood protein vitronectin bound to lipid and HDL"

**Table S1. System for MD simulation.** Five independent simulations were initiated form from the coordinates of $Ca^{2+}$-bound Vn-HX (PDB 7rj9; Tian et al. *Biophys. J.* **2022,** 121, 3896-3906).

| | | | Number of Molecules | | | |
|---|---|---|---|---|---|---|
| **System** | **System size (Å and angles)** | **Total MD simulation time** | **VnHX** | **Lipids and ApoA1** | **Ions** | **Water** |
| **VnHX–diC7PC** | 120.00 x 120.00 x 120.00 Å<br>90 x 90 x 90°<br>(cube) | 6 x 6 μs = 36 μs | 1 | diC7-PC: 20 | $Cl^-$: 302<br>$Na^+$: 296<br>$Ca^{2+}$: 2 | 53,041 |
| **VnHX–POPC$_{s1}$** | 117.37 x 117.37 x 117.37 Å<br>90 x 90 x 90°<br>(cube) | 2 x 4.5 μs = 9 μs | 1 | POPC: 1 | $Cl^-$: 302<br>$Na^+$: 296<br>$Ca^{2+}$: 2 | 53,041 |
| **VnHX–POPC$_{s2}$** | 117.37 x 117.37 x 117.37 Å<br>90 x 90 x 90°<br>(cube) | 2 x 4.5 μs = 9 μs | 1 | POPC: 1 | $Cl^-$: 302<br>$Na^+$: 296<br>$Ca^{2+}$: 2 | 53,041 |
| **VnHX–POPC$_{s1s2}$** | 117.40 x 117.40 x 117.40 Å<br>90 x 90 x 90°<br>(cube) | 2 x 4.5 μs = 9 μs | 1 | POPC: 2 | $Cl^-$: 302<br>$Na^+$: 296<br>$Ca^{2+}$: 2 | 53,041 |
| **VnHX–LPP$_{10μs}$** | 183.94 x 183.94 x 183.94 Å<br>60 x 60 x 90°<br>(rhombic dodecahedron) | 3 x 5 μs = 15 μs | 1 | POPC: 161<br>Cholesterol: 24<br>ApoA1: 2 | $Cl^-$: 427<br>$Na^+$: 431<br>$Ca^{2+}$: 1 | 137878 |
| **VnHX–LPP$_{20μs}$** | 191.04 x 191.04 x 191.04 Å<br>60 x 60 x 90°<br>(rhombic dodecahedron) | 3 x 5 μs = 15 μs | 1 | POPC: 161<br>Cholesterol: 24<br>ApoA1: 2 | $Cl^-$: 468<br>$Na^+$: 472<br>$Ca^{2+}$: 1 | 153592 |

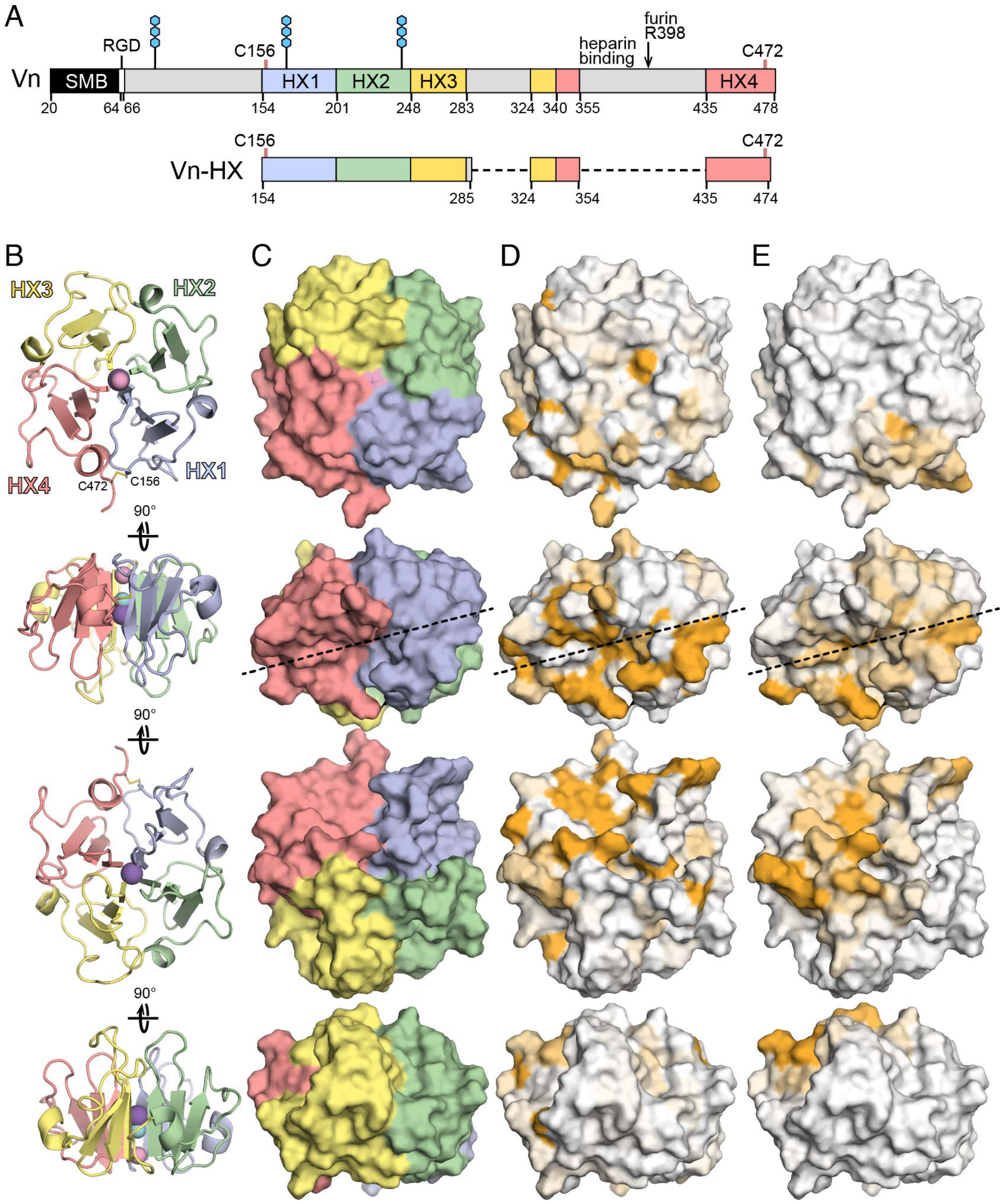


**Figure S1. Vn sequence and structure.**

**(A)** Vn domain organization. Mature Vn starts at Asp20 and comprises a somatomedin B (SMB) domain (black), an ArgGlyAsp (RGD) sequence (white), an HX domain colored by repeat unit (HX1, blue; HX2, green; HX3, yellow; HX4, red), and three regions (gray) with lower sequence conservation and higher predicted disorder. C156 and C472 form a disulfide bond. Blue hexagons denote N-glycosylation sites.

**(B-E)** Structure of the Vn-HX domain, cartoon and surface representations. Colors reflect HX repeat units (B, C), lipid-induced NMR perturbations (D), or frequency of contacts with lipid moieties from MD simulations (E). The dashed line marks the hydrophobic cleft. The propeller is circularized by the C156-C472 disulfide bond. Each propeller blade is formed by three $\beta$ strands ($\beta$1-$\beta$3). Spheres denote occluded $Na^+$ (slate), $Cl^-$ (aqua) and $Ca^{2+}$ (pink) ions. Shown for molecule A of the asymmetric unit; PDB 7rj9.

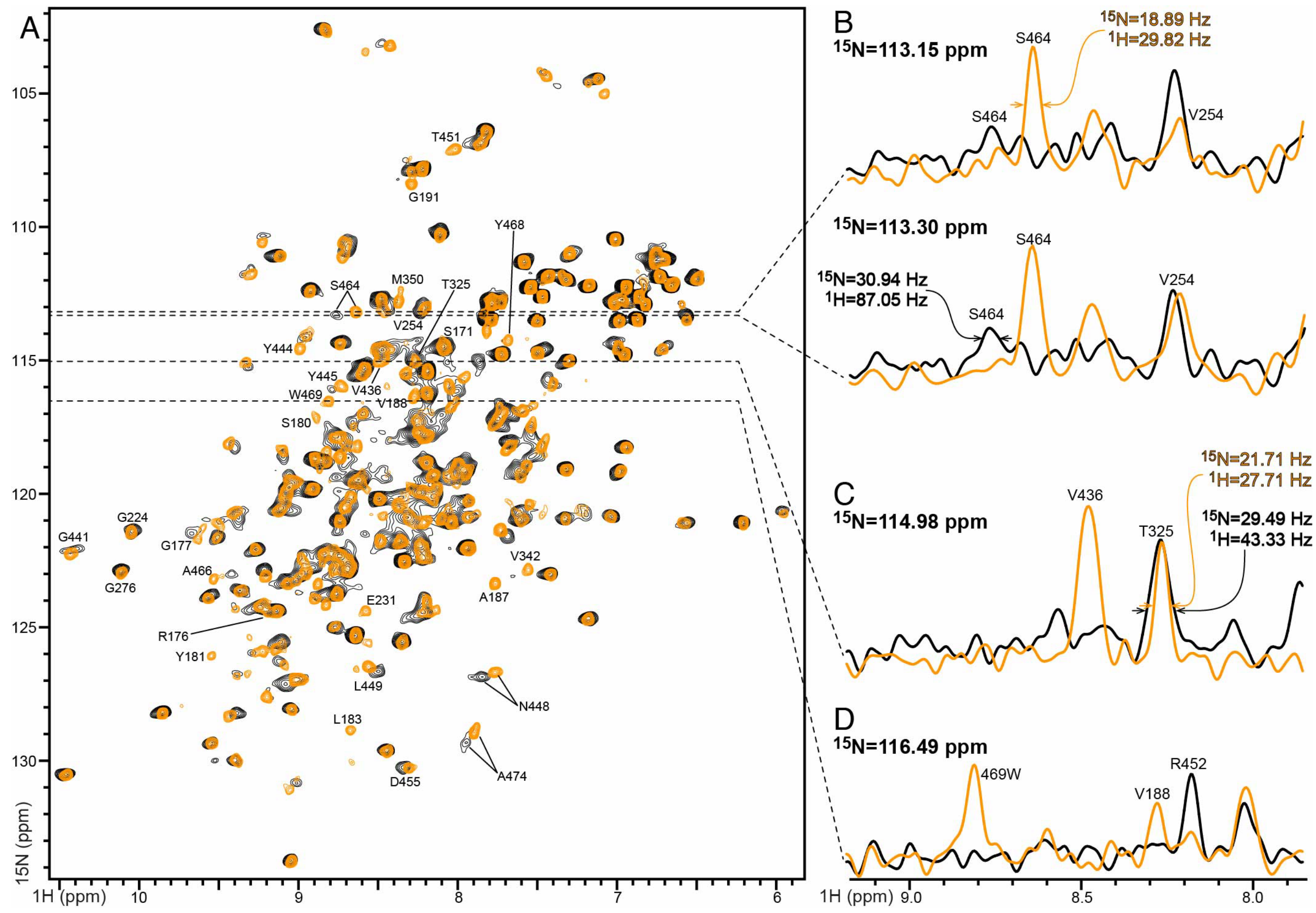


**Figure S2. $^{1}$H/$^{15}$N NMR spectra of $^{15}$N labeled Vn-HX without (black) or with (yellow) diC7-PC. (A)** 2D $^{1}$H/$^{15}$N HSQC spectra. **(B-D)** 1D $^{1}$H spectral slices from the 2D spectrum. The molar ratio of diC7-PC to protein is 50/1. Examples of new peaks (A, C, D), chemical shit perturbations (A, B), and line narrowing (D) observed upon addition of diC7-PC are marked.

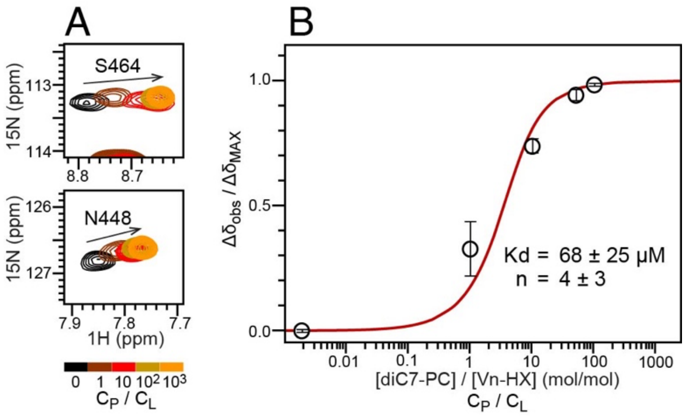


**Figure S3. Lipid-binding affinity derived from diC7-PC induced changes in solution NMR HSQC $^{1}H/^{15}N$ chemical shifts. (A)** Representative $^{1}H/^{15}N$ NMR signals of Vn-HX with increasing concentrations of diC7-PC (black to yellow) from 0 to $10^3$ lipid/protein ($C_P/C_L$) molar ratio. **(B)** Change in $^{1}H/^{15}N$ NMR chemical shift as a function of lipid/protein molar ratio. Each data point is the average value of eight well-resolved $^{1}H/^{15}N$ NMR signals with mean standard error. Values of the apparent dissociation constant (Kd) were estimated by fitting the data for each $^{1}H/^{15}N$ signal to an n-site binding model described by the equation:

$$\Delta\delta_{obs} / \Delta\delta_{MAX} = \{ (nC_P+C_L+K_d) - [ (nC_P+C_L+K_d)^2 - 4nC_PC_L) ]^{1/2} \} / (2nC_P)$$

where $\Delta\delta_{obs}$ is the observed change in chemical shift, $\Delta\delta_{MAX}$ is the maximum change observed at saturation, n is the number of binding sites, $C_P$ is the protein concentration and $C_L$ is the lipid concentration. The best fits yield an average Kd of 68±25 μM and n=4 (red line).

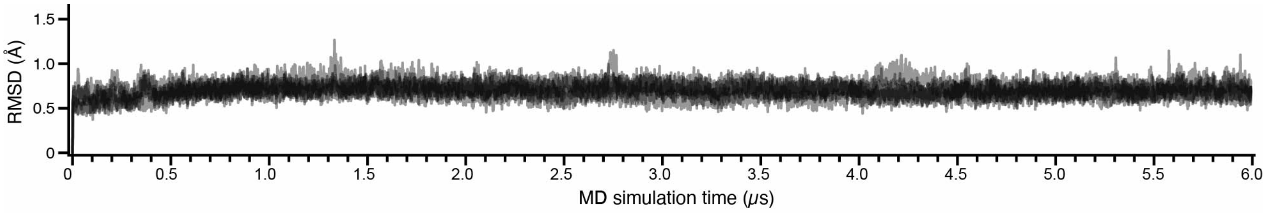


**Figure S4. MD simulations of Vn-HX in diC7-PC.** Time evolution of protein coordinates, including only the β-propeller proper and excluding flexible regions. Each trace represents one of five independent MD simulations.

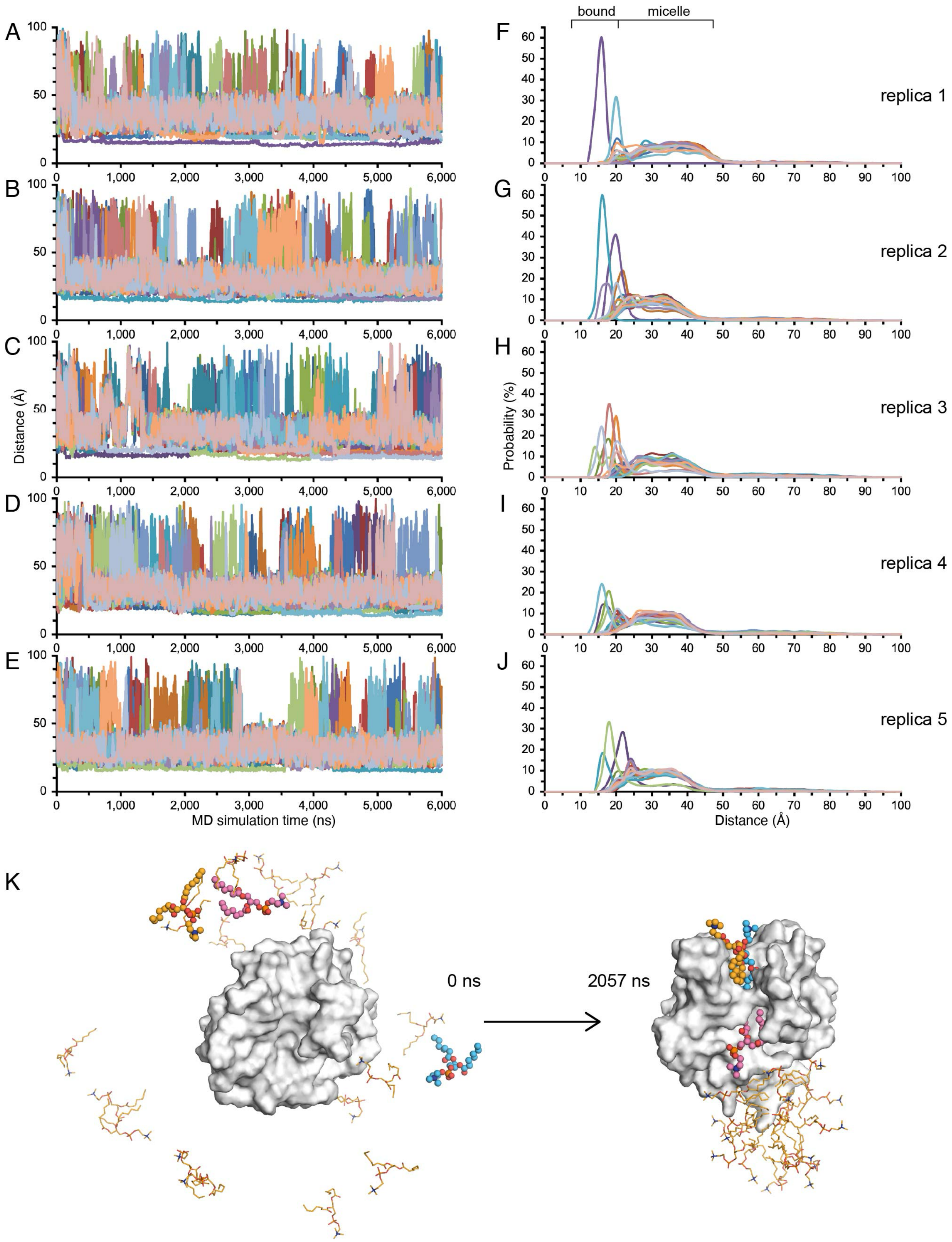


**Figure S5. MD simulations of Vn-HX in diC7-PC. (A-J)** Time evolution (A-E) and probability distribution (F-J) of the center of mass distance between Vn-HX and each diC7-PC lipid molecule. Each trace represents one of twenty lipid molecules in the simulation. **(K)** Representative MD simulation of lipid binding from 0 ns to 2057 ns.

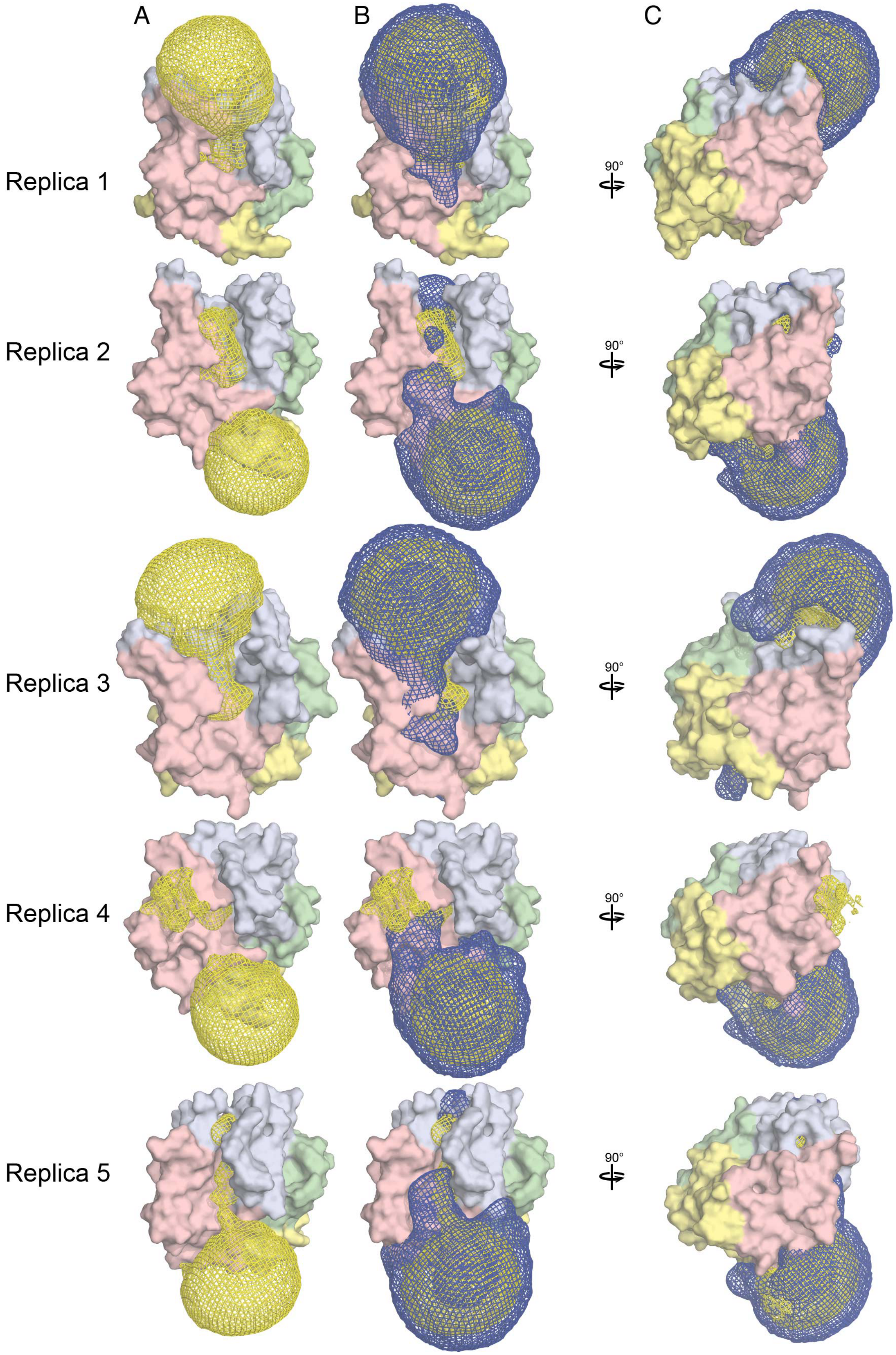


**Figure S6. Volumetric map analysis.** Volume densities (mesh representation) represent the time-average for each of the five replicas. They are shown for the lipid polar headgroups (blue mesh) or lipid acyl chains (yellow mesh) individually (A) or together (B, C).

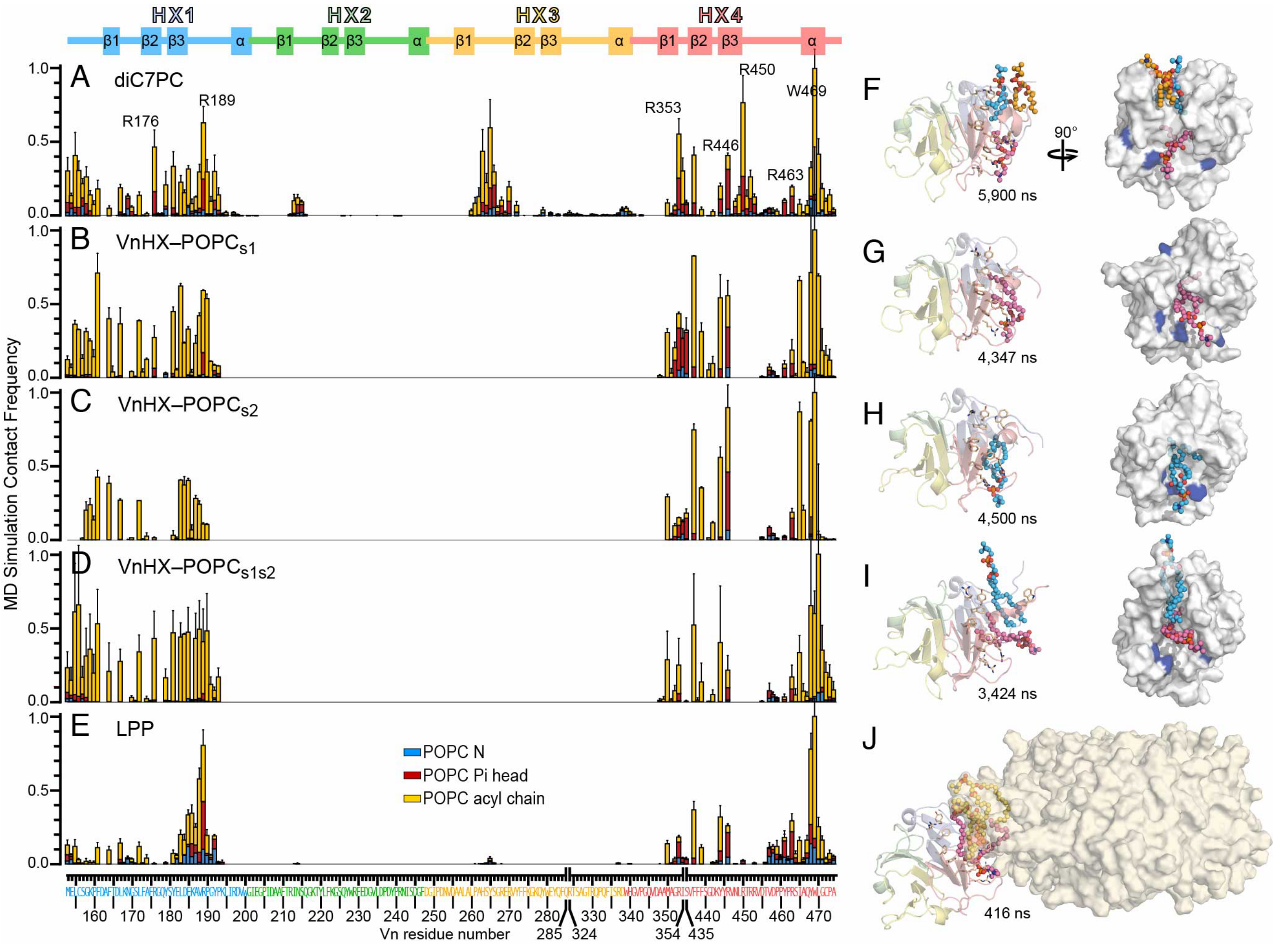


**Figure S7. MD simulations map the lipid binding site of Vn.** Protein-lipid contact frequency from MD simulations. Data are averaged over all replicas. The bars represent the frequency of finding a heavy atom of each protein residue within 4Å of heavy atoms in the lipid acyl chain (yellow), or polar headgroup phosphate (red) and trimethylammonium nitrogen (blue). Error bars denote standard error of the mean. **(F-J)** Representative structures, taken at specific time points of MD simulation. Spheres (pink, blue, yellow) denote lipid bound at site 1 (pink), site 2 (blue) or recruited (yellow) from the pool of diC7PC (F) or the lipoparticle (J). VnHX color reflects repeat units (HX1, blue; HX2, green; HX3, yellow; HX4, red). Blue surface denotes key gate Arg. The HDL lipoparticle (beige Connolly surface) is shown without ApoA1 for simplicity (J).

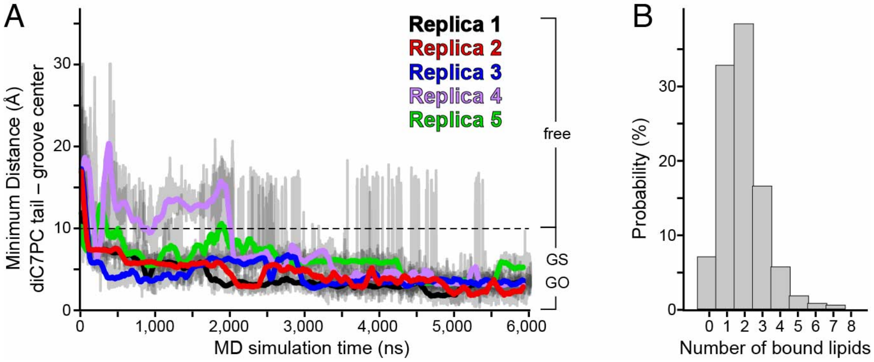


**Figure S8. Lipid binding site of Vn. (A)** Time evolution of the minimum distance between the lipid tails and hydrophobic groove. Each colored trace is the running average over a period of 100 ns for each replica of MD simulation. Gray traces are the full data for each replica. The minimum distances reflect lipid bound to the groove surface (GS), or occluded within its aromatic cage (GO). **(B)** Probability of locating lipids within 10 Å of the groove core center.

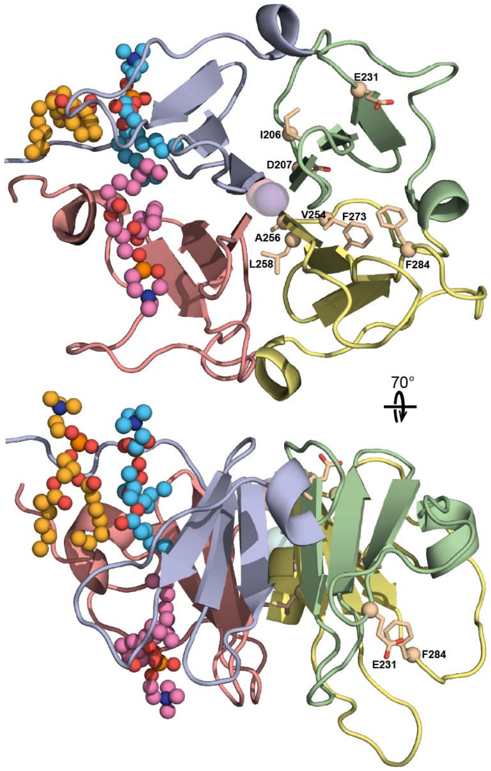


**Figure S9. Sites with lipid-perturbed NMR signals but no MD atomic contacts.** Representative structure, taken at 2,057 ns of MD simulation, with lipid molecules (pink, blue, yellow spheres) bound to the HX1-HX4 groove, and key residues (wheat sticks) with $^{1}H/^{15}N$ signals perturbations but no MD atomic contacts with lipid. Colors (pink, blue, yellow) identify each bound lipid molecule.

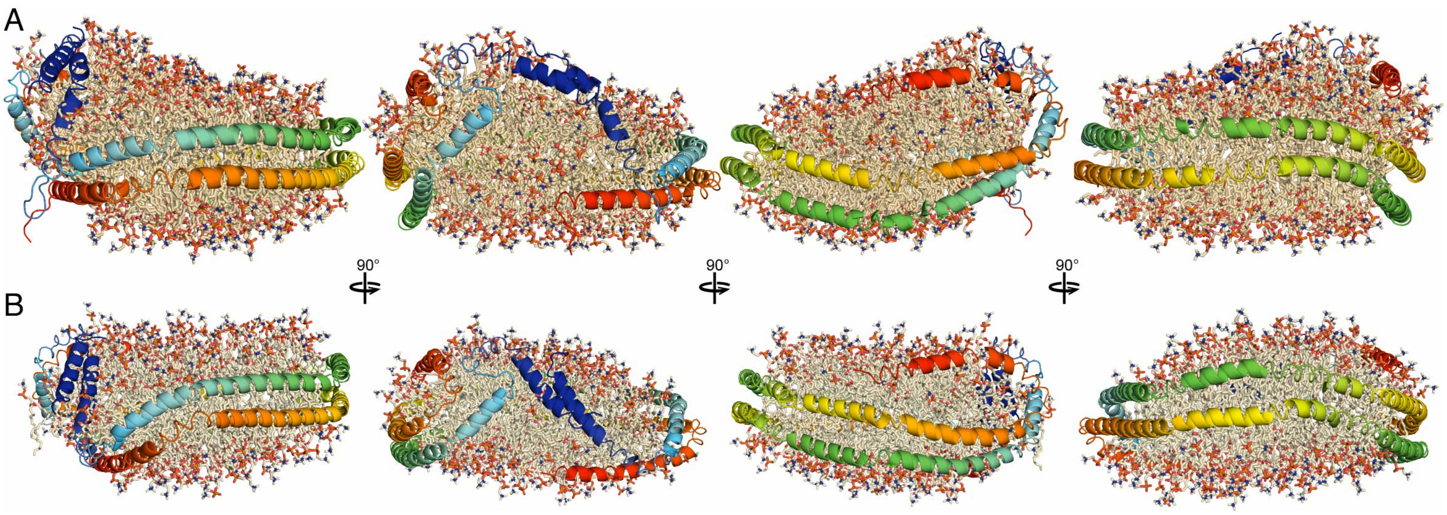


**Figure S10. Starting models of HDL for MD simulation. (A, B)** Orthogonal views of the structural models taken at 10 μs (A) or 20 μs (B) of the published MD simulation trajectory of a 100 Å diameter HDL lipoparticle (Pourmousa et al. 2018 *Proc. Natl. Acad. Sci. U.S.A.* 115: 5163). Lipids and cholesterol are shown as sticks. ApoA1 is shown as cartoon in blue-red color gradient to denote N to C terminus progression.

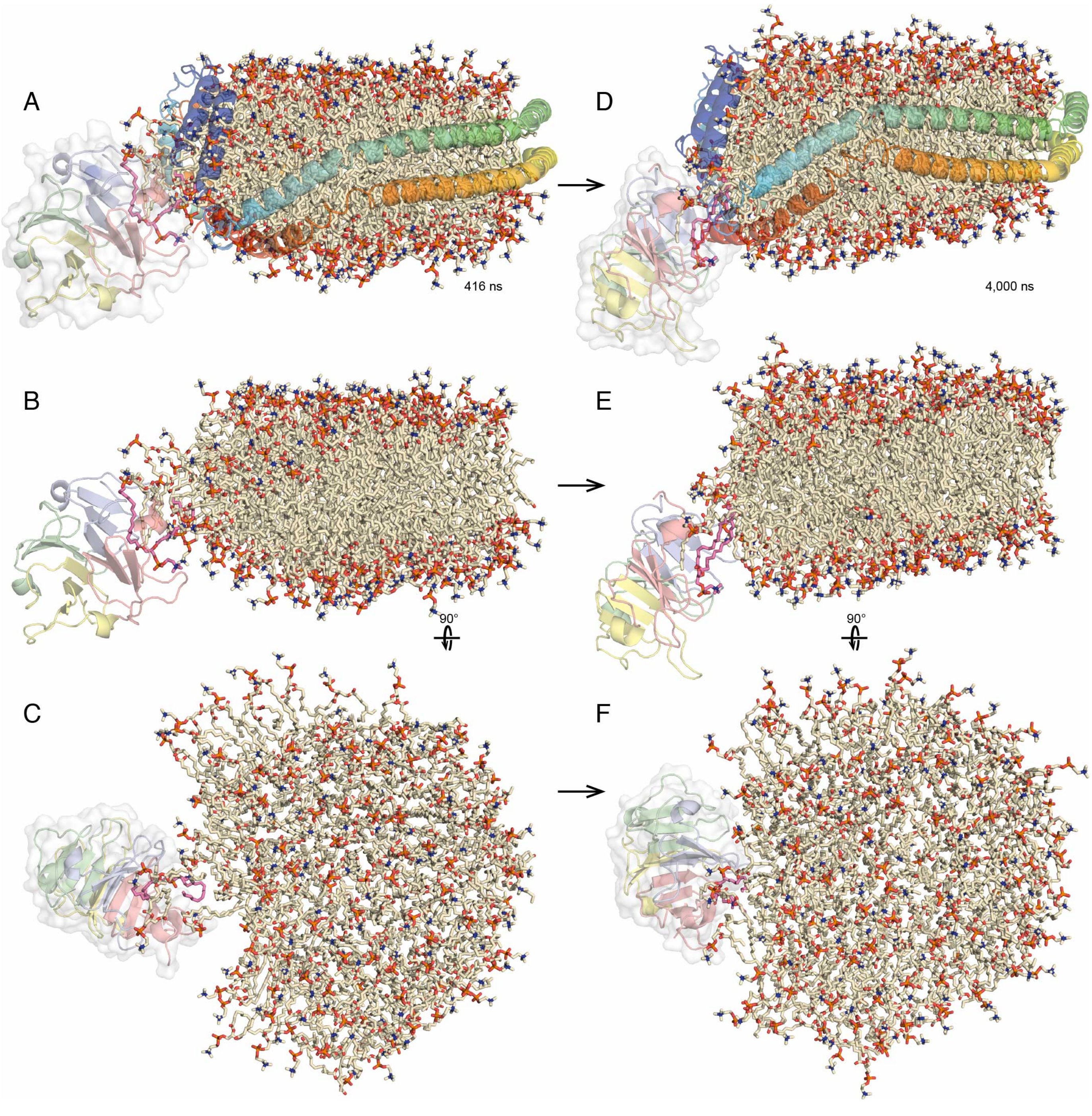


**Figure S11. VnHX–HDL association.** Time evolution of the structural models from 416 ns (A-C) to 4,000 ns (D-F. ApoA1 is shown as cartoon (A, D) or removed from the image (B, C, E, F) to show the lipids. ApoA1 blue-red color gradient denotes N to C terminus progression. POPC color denotes the original $POPC_{s1}$ (pink), or $POPC_{HDL}$ pool (wheat).
